# Flagellar motor remodeling during swarming requires FliL

**DOI:** 10.1101/2023.07.14.549092

**Authors:** Jonathan D. Partridge, Yann Dufour, YuneSahng Hwang, Rasika M. Harshey

## Abstract

FliL is an essential component of the flagellar machinery in some bacteria, but a conditional one in others. The conditional role is for optimal swarming in some bacteria. During swarming, physical forces associated with movement on a surface are expected to exert a higher load on the flagellum, requiring more motor torque to move. Bacterial physiology and morphology are also altered during swarming to cope with the challenges of surface navigation. FliL was reported to enhance motor output in several bacteria and observed to assemble as a ring around ion-conducting stators that power the motor. In this study we identify a common new function for FliL in diverse bacteria – *Escherichia coli, Bacillus subtilis* and *Proteus mirabilis*. During swarming, all these bacteria show increased cell speed and a skewed motor bias that suppresses cell tumbling. We demonstrate that these altered motor parameters, or ‘motor remodeling’, require FliL. Both swarming and motor remodeling can be restored in an *E. coli fliL* mutant by complementation with *fliL* genes from *P. mirabilis* and *B. subtilis*, showing conservation of swarming-associated FliL function across phyla. In addition, we demonstrate that the strong interaction we reported earlier between FliL and the flagellar MS-ring protein FliF is confined to the RBM-3 domain of FliF that links the periplasmic rod to the cytoplasmic C-ring. This interaction may explain several phenotypes associated with the absence of FliL.

## INTRODUCTION

Bacterial flagella are vital organelles that facilitate the survival and spread of bacteria in a range of environments (Harshey, 2003, Jarrell and McBride, 2008, Duan et al., 2013, Chaban et al., 2015, Wadhwa and Berg, 2022). In model organisms *E. coli* and *Salmonella*, peritrichous flagella are anchored within the cell membranes via many connecting segments. The long external filament is attached via a flexible external hook to a rod that traverses the periplasm, attaching to a basal MS ring (FliF) in the inner membrane (Chevance and Hughes, 2008, Johnson et al., 2021, Tan et al., 2021). L– and P-rings in the outer membrane act as a bushing for the rotating rod. The extreme C– terminus of FliF interacts with FliG atop the cytoplasmic C ring. The MS and C rings together are referred to as the rotor. The rotor is surrounded by a varying number of proton-conducting stator units (MotA5::MotB2 complex) (Deme et al., 2020, Santiveri et al., 2020, Tan et al., 2021). The rotor and stators together are called the motor (Berg, 2003). The stator complexes are dynamic, moving in an out of the rotor by sensing how much torque is needed to drive the rotor (Tipping et al., 2013, Lele et al., 2013). When drifting in the membrane, ion flow through the stators is shut off by a ‘plug’ contributed by the MotB subunit of complex, opening only when the stators dock at the rotor, and when MotA engages with FliG in the C ring (Kojima et al., 2009, Hosking et al., 2006). Ion flow through the stators changes the charge of FliG residues at the MotA-FliG interface, producing torque and rotor movement (Berg and Turner, 1993, Zhou et al., 1998). While a single stator is sufficient to drive rotation, up to 11 stators can maximally assemble around the rotor (Reid et al., 2006). Stator numbers are a general reflection of load exerted upon the flagellar filament.

The rotor is bi-directional, with counter-clockwise rotation (CCW) producing flagella bundling that results in a run or forward motion of the cell, and clock-wise rotation (CW) producing flagella unbundling that results in a tumble, randomizing cell reorientation (tumble) (Berg, 2003). The default state of the rotor is CCW. Bidirectionality is imposed by a chemosensory system that detects attractant and repellent environmental stimuli to control the levels of the signaling protein CheY∼P, which interacts with the C-ring to produce the CW intervals. By biasing the frequency of CW events, cells can decrease or increase their tumble bias (TB; defined as the time spent tumbling over the total trajectory time of each cell) as they drift towards attractants or away from repellents.

Flagella also propel bacteria on a surface in a movement called swarming, where bacteria move as a collective (Harshey, 2003, Be’er and Ariel, 2019, Kearns, 2010, Partridge, 2022, Darnton et al., 2010, Partridge and Harshey, 2013b). The surface presents several challenges to movement, some of these being inadequate liquid for the flagella to push through, as well as physical forces/properties such as surface tension/adhesion/friction/viscosity (Tuson and Weibel, 2013). Bacteria overcome these obstacles in different ways, some by secreting osmolytes or surfactants, others by changing cell shape and/or increasing flagella production, and still others by employing different sets of stators (Baker and O’Toole, 2017). That it takes increased motor torque to navigate a surface is attested to by the requirement for increased flagella numbers (Ghelardi et al., 2012, McCarter et al., 1988, Mukherjee et al., 2015), special stators (Toutain et al., 2005), and stator-associated proteins such as FliL or SwrD during swarming in different bacteria (Hall et al., 2018, Attmannspacher et al., 2008). FliL is also essential for swimming motility of polarly flagellated *Caulobacter crescentus* (Jenal et al., 1994) and *Rhodobacter sphaeroides* (Suaste-Olmos et al., 2010), and for lateral flagella-driven swimming motility of *Bradyrhizobium diazoefficiens* (Mengucci et al., 2020).

A variety of bacteria alter motor behavior during swarming, increasing motor speed and suppressing tumbles by altering motor bias (Partridge et al., 2019, Partridge et al., 2020, Tian et al., 2021). We refer to both effects as ‘motor remodeling’. The resulting increased time spent in forward motion likely contributes to cell alignment and cohesion of the collectively moving swarm. The present study was initiated not only because FliL has a profoundly negative effect on swarming in *E. coli* and *Salmonella* (Attmannspacher et al., 2008), but also because it affects motor bias and speed (the motor properties remodeled during swarming) even in planktonic bacteria, as monitored by single motor analysis (Partridge et al., 2015). According to secondary structure predictions, FliL traverses the inner membrane once in these bacteria, with the bulk of the protein residing in the periplasm (Attmannspacher et al., 2008). Strong FliL-FliL, FliL-MotA, FliL-MotB and FliL-FliF interactions in pull-down and 2-hybrid assays suggested that FliL is positioned near the stators (Partridge et al., 2015). This assignment was corroborated by cryo-ET and structural studies of FliL in other bacteria, which found that FliL forms a decameric ring around the stators (Takekawa et al., 2019, Tachiyama et al., 2022, Guo et al., 2022). Based on a variety of studies in different bacteria, a consensus has emerged that FliL is required to produce more torque needed for rotating flagella, which would be the case when moving over a surface or through viscous media (Partridge et al., 2015, Zhu et al., 2015, Lin et al., 2018, Homma and Kojima, 2022, Tachiyama et al., 2022, Guo and Liu, 2022, Guo et al., 2022). Besides a functional role, a structural role for FliL at the base of the flagellum has been inferred in *E. coli* and *Salmonella*, where absence of FliL leads to rod fracture only during swarming (Attmannspacher et al., 2008, Partridge et al., 2015). In *Borrelia burgdorferi*, absence of FliL produced an abnormal orientation of the periplasmic flagella, indicating a structural contribution here as well (Motaleb et al., 2011).

We show in this study that motor remodeling requires FliL not only in *E. coli*, but also in *B. subtilis* and *P. mirabilis*. Surprisingly, we find that FliL from the latter two bacteria can function in *E. coli*. Pull-down experiments show a strong interaction of *E. coli* FliL with the major D3 domain of the MS ring protein FliF, which may help explain the many FliL phenotypes.

## RESULTS

### Cell tracking experiments reveal that absence of FliL abrogates remodeling of the motor during swarming

*E. coli* cells inoculated on standard plates set with 0.5% Eiken agar do not swarm in the absence of FliL, and a subset of cells lifted from the agar show rod fracture (Attmannspacher et al., 2008). To track the motion of cells whose flagella were still intact, we lifted them from *fliL* cells propagated on a swarm plate, transferred them into liquid, and first ascertained that sufficient numbers were still motile (Fig. S1; see also supplementary videos). Trajectories of the motile cells were monitored in a pseudo-two-dimensional channel for 100 s and analyzed as reported earlier (Partridge et al., 2019). Four sets of cells/conditions were compared: WT and *fliL*, taken from either liquid or swarm plates. In liquid, WT cells had a median speed of 22.6 µm/s while that of *fliL* cells was 15.85 µm/s, the 30% drop in *fliL* speed consistent with earlier reports using single motors (Partridge et al., 2015) (Fig. 1A) (statistics are detailed in Table S1). In a swarm, WT cells increase speed compared to liquid conditions as reported previously (Partridge et al., 2019). While *fliL* cells also increased speed, the values were lower than WT (25.4 µm/s vs 18.51 µm/s). The median tumble bias (TB) of WT was 0.11 in liquid, dropping to 0.07 in the swarm, while that of the *fliL* mutant was higher than the WT in liquid (0.16), and increased further to 0.26 in the swarm (statistics in Table S1). Thus, the *fliL* motor fails to remodel on swarm agar.

**Figure 1:**
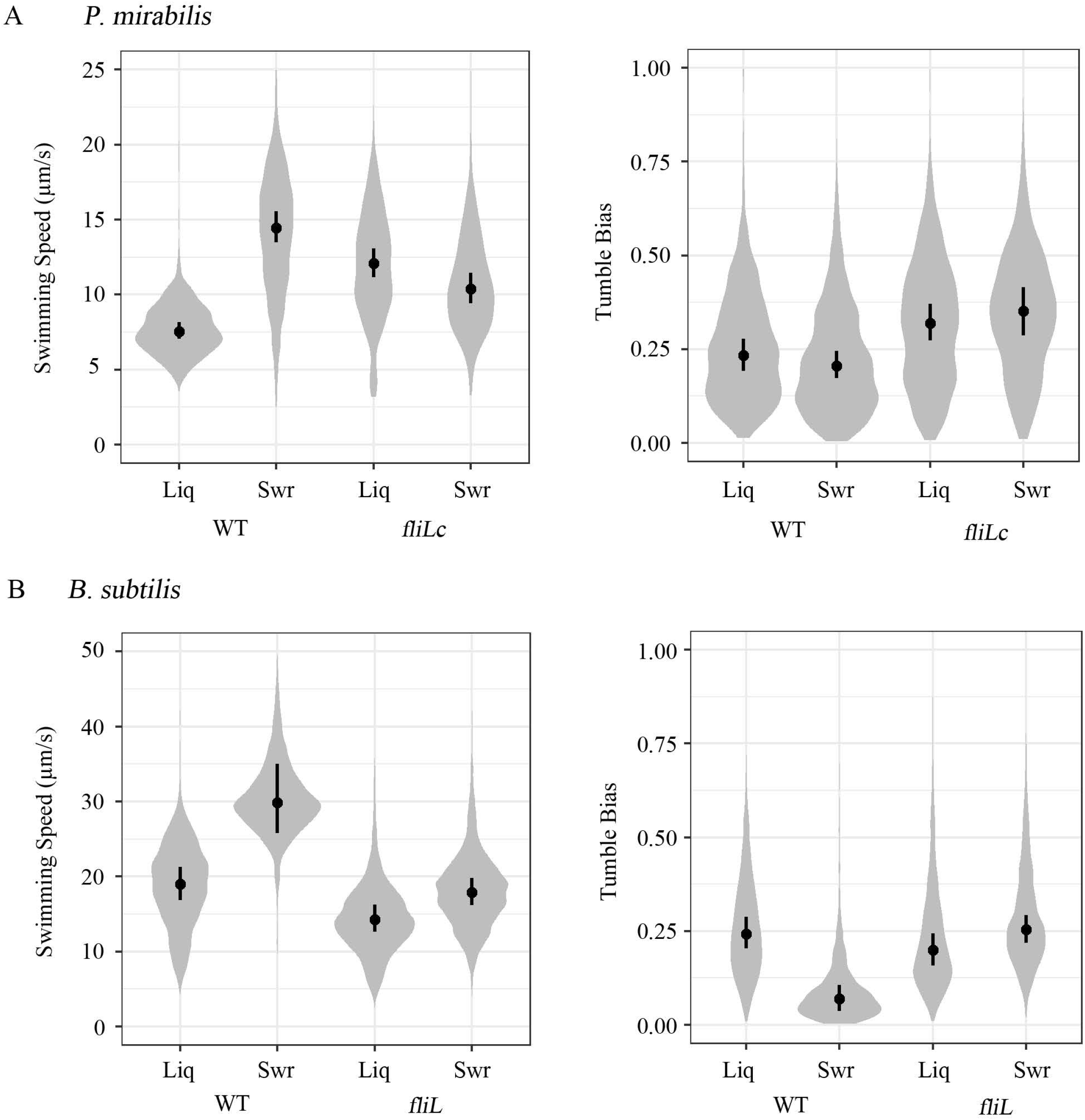
Motor behavior of *E. coli* WT and *fliL* cells in liquid and swarming cultures. A) Probability density distribution of swimming speeds (left) and cell tumble biases (right) within a pseudo-2D environment. Cells were grown in LB (Liq) or LB swarm agar (Swr) before transfer to LB liquid for observation (see Methods). Cell movement was recorded for 100 s using phase-contrast microscopy at 10x magnification. Each distribution was calculated from more than 10,000 individual trajectories (>2,300 min of cumulative time) for each condition, combined from five independent experiments for each strain/condition. Black dots and bars indicate the median and 95% credible intervals of the posterior probabilities of the medians for each treatment. B) Behavior of single motors of *E. coli* WT and *fliL* cells over a 60-s time frame. Rotation was monitored by recording the motion of 0.75 µm polystyrene beads attached to sheared “sticky” filaments. Rotation speeds (left) are expressed in Hz, and Switching frequency as reversals/min (middle). Torque (piconewtons per nanometer) was derived from CCW motor speeds (right). At least 15 motors were recorded for each condition. Black bars indicate the mean value, calculated *p* values indicate statistical significance (<0.005) for comparisons of strains (WT vs *fliL*) and conditions (Liq vs Swr) for both speed and TB values (Table S1).

The various physical forces that bacteria must overcome in order to swarm on a surface (wetness/ tension/ adhesion/ friction/ viscosity) are expected to vary depending on the agar concentration i.e. how ‘soft’ or ‘hard’ its texture is. Indeed, bacteria have been classified into ‘temperate’ or ‘robust’ swarmers on the basis of this criterion (Partridge and Harshey, 2013b). This classification is likely due to how much motor power a particular species is able to generate, or how much water is available on the surface, or how much surfactant it makes. For example, *Salmonella*, which will normally swarm on media solidified with 0.5%-0.6% w/v agar, could swarm on media with up to 1% w/v agar when motor output was increased by overproducing either flagella or FliL plus MotAB (Partridge and Harshey, 2013a). *E. coli* is even fastidious about the commercial source of the agar (Eiken; (Harshey and Matsuyama, 1994)), which has undefined properties that are ‘permissive’ for *E. coli* to swarm. Indeed, *Salmonella* was able to swarm more efficiently on Eiken agar compared to Fisher agar (Fig. S2A, left). If the *fliL* swarming defect was related to its lower motor torque, we wondered if lowering the agar concentration further or hydrating the surface would compensate: both manipulations partially rescued swarming (Fig. S2B; only hydration experiment shown). Similarly, the permissive Eiken agar partially rescued the swarming defect of a *Salmonella fliL* mutant (Fig. S2A, right).

*E. coli* WT and *fliL* cells were tracked again under these more permissive swarming conditions. For WT, hydration increased speed further from 25.4 µm/s to 27.2 µm/s, while TB dropped from 0.07 to 0.04 (Fig. S2C) (statistics in Table S1). For *fliL*, hydration did not increase speed, which dropped from 18.5 to 16.9 µm/s, but TB decreased from 0.27 to 0.13. We do not understand the latter result, but note that the reduced TB of this mutant is still much higher than that observed in WT cells (Fig. S2C).

Taken together, the data in Fig. 1A and Fig. S2 corroborate the contribution of FliL to maintaining a high swimming speed and reveal in addition a FliL contribution to reducing TB during swarming. Partial rescue of the *fliL* swarming defect by a ‘wetter’ agar surface suggests that less motor power is required under these conditions.

### Single motor analysis corroborates cell tracking data for *fliL* cells

To corroborate the results obtained from cell tracking we monitored the output of individual motors using the bead assay, as previously described (Partridge et al., 2015, Partridge et al., 2019). In this assay, a polystyrene bead is attached to a sheared filament stub, and its rotation speed and bias recorded by a high-speed camera (Terasawa et al., 2011, Ryu et al., 2000).

WT *E. coli* cultivated in liquid media showed average single motor angular velocity of 70.1 Hz ±5.8 Hz, increasing to 82.92 Hz ±7.29 Hz in the swarm, while motor reversals dropped significantly from 38 ±5 reversals per min (rv.p.m.) in liquid to 8.6 ±2.8 rv.p.m in the swarm (Fig. 1B), values similar to those that we reported previously (Partridge et al., 2019). Rv.p.m refers to the frequency of shifts from CCW (the basal state of the motor) to CW rotation. Under these conditions, *fliL* motors showed lower speeds (58.6 Hz ±9.2) and higher reversals (27.1 rv.p.m ±4.5/ 23.2 rv.p.m ±4.2, liquid / swarm) under both growth conditions. The torque of the WT flagellar motor increased from 648.1 ±27 pN in liquid to 702.7 ±27.3 pN under swarm conditions, in line with previous observations (Partridge et al., 2019). The *fliL* motor did not behave similarly (Partridge et al., 2015), showing half the torque of the WT motor in both media: 345 ±27.8 pN in liquid, and 358.7 ±25.3 pN from swarms.

In summary, the speed and bias parameters derived from individual motors are in overall agreement with those from cell tracking (Fig. 1A). A prominent effect on motor remodeling during swarming is the drop in switching frequency, which the *fliL* motor is unable to execute.

### FliL effects on motor remodeling are seen in diverse bacteria

Motor remodeling associated with swarming is widespread in swarming bacteria (Partridge et al., 2020). To test if loss of FliL affects remodeling in these other bacteria as shown above for *E. coli*, we chose two bacteria – *B. subtilis* and *P. mirabilis* – where an effect of FliL on swarming has been demonstrated (Lee and Belas, 2015, Cusick et al., 2012, Hall et al., 2018). We obtained a *fliL* knockout mutant of *B. subtilis* from Daniel Kearns (Indiana University), which exhibits only a partial swarming defect (Hall et al., 2018). We were unable to either obtain or generate a similar mutant reported in *P. mirabilis*. We therefore acquired a polar Tn5 mutant of *fliL* in *P. mirabilis* from Phillip Rather (Emory University), which we used to isolate motile suppressors as reported earlier (Cusick et al., 2012). The suppressor we isolated was identical to that previously reported by Cusick et al. (YL1001), where partial excision of Tn5 altered 28 residues at its C terminus; we will refer to this variant as *fliLC*. The solved structures of FliL show that the C-terminus forms a conserved β strand (Takekawa et al., 2019, Tachiyama et al., 2022), absence of which would likely either destabilize or alter the fold of the periplasmic domain of FliL (Anna Roujeinikova, personal communication). This suppressor was reported to swarm (Cusick et al., 2012, Guo et al., 2022). Unlike the *E. coli fliL* null, neither the *B. subtilis fliL* null nor the *P. mirabilis fliLC* mutants abolished swarming. As reported, the *B. subtilis* mutant showed reduced swarming at both 0.5% and 0.7% agar compared to WT, while the *P. mirabilis* showed a smaller swarm colony diameter only at 0.7% and 1% agar (Fig. S3). For tracking, cells were lifted from 0.5% Eiken agar plates as done for *E. coli*. Motility was monitored for both WT and *fliL* bacteria as before. For *B. subtilis*, we used the surfactin-versions of the strains because surfactin interferes with tracking (Partridge et al., 2020).

Motility parameters for WT *P. mirabilis* aligned with previous observations (Partridge et al., 2020), with speed increasing and TB dropping as cells transitioned from liquid to swarm conditions (Fig. 2A; see Table S1). The *P. mirabilis fliLC* was faster than its WT counterpart in liquid, but not in the swarm, and had an increased TB in both liquid and swarm (Fig. 2A). Motility parameters for *B. subtilis* WT also aligned with those reported earlier (Partridge et al., 2020), with speeds increasing and TBs dropping in the swarm (Fig 2B, Table S1). The *B. subtilis fliL* mutant affected both liquid– and swarm speeds and failed to decrease TB in the swarm (Fig. 2B).

**Figure 2:**
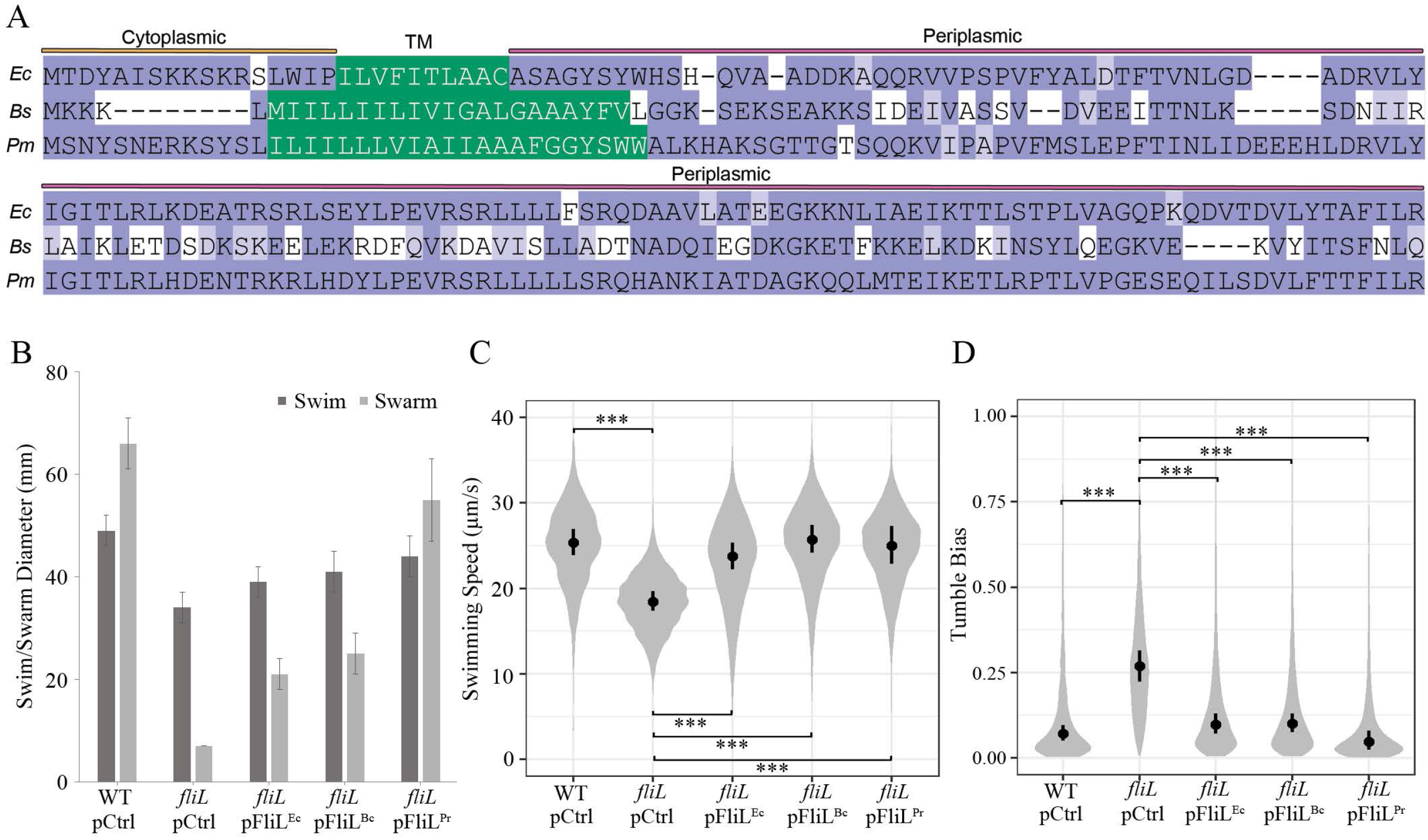
Tracking *B. subtilis* and *P. mirabilis* WT and *fliL* cells. Swimming speed (left) and tumble bias (right) probability density distributions for A) *P. mirabilis* (JP2891) and B) *B. subtilis* (DS191). Cells were grown in LB (Liq) or LB swarm agar (Swr) with glucose (0.5% w/v), before transfer to LB liquid for observation. Each distribution was calculated from more than 6,500 individual trajectories (>800 min of cumulative time) for each condition, combined from at least three independent experiments for each strain/condition. Black dot and bars indicate the median and 95% credible intervals of the posterior probabilities of the medians for each treatment. See Table S1 for *p* values.

We conclude that a complete absence of FliL in *B. subtili*s or an altered FliL C-terminus in *P. mirabilis*, both affect motor remodeling as seen in *E. coli fliL*.

### *E. coli fliL* mutant can be rescued with FliL from *P. mirabilis* and *B. subtilis*

FliL sequences from the three bacteria used in this work are shown in Figure 3A, and their predicted Alpha-fold structures in Fig. S4A. Given the similarities in their structure, and similar contribution of all three proteins to motor behavior in their respective hosts, we wondered if their function might be interchangeable between the three species. The cell envelope in which the flagellum is embedded is very different between Gram negative vs Gram positive bacteria, with absence of a periplasm in the latter (Mukherjee and Kearns, 2014). Surprisingly, plasmids expressing *fliL* from *Bacillus* (FliL^Bc^), and *Proteus* (FliL^Pr^), complemented an *E. coli fliL* null mutant for both swimming and swarming (Fig. 3B, Fig. S4B). We note that a plasmid expressing *E. coli fliL* (FliL^Ec^) did not achieve full complementation. Perhaps incorporation of FliL at the basal body depends on the order of gene expression from the *fliLMNOPR* operon, ectopic expression being untimely. Alternatively, the stoichiometry of FliL matters. Surprisingly however, FliL^Pr^ was more proficient at rescuing the *E. coli fliL* defect than FliL^Ec^, giving nearly WT levels of motility. We cross-checked this result using a *Salmonella fliL* mutant (FliL^Se^; Fig. S4C). Here again, we observed robust complementation of the *Salmonella fliL* defect with FliL^Pr^. While FliL^Se^ complementation was suboptimal, neither FliL^Ec^ nor FliL^Bc^ complemented *Salmonella fliL* for swarming. The negative results for this set of cross-species complementation results reflect only the difficulty of FliL complementation, given that complementation of *fliL* defects with FliL from the same species is suboptimal.

**Figure 3:**
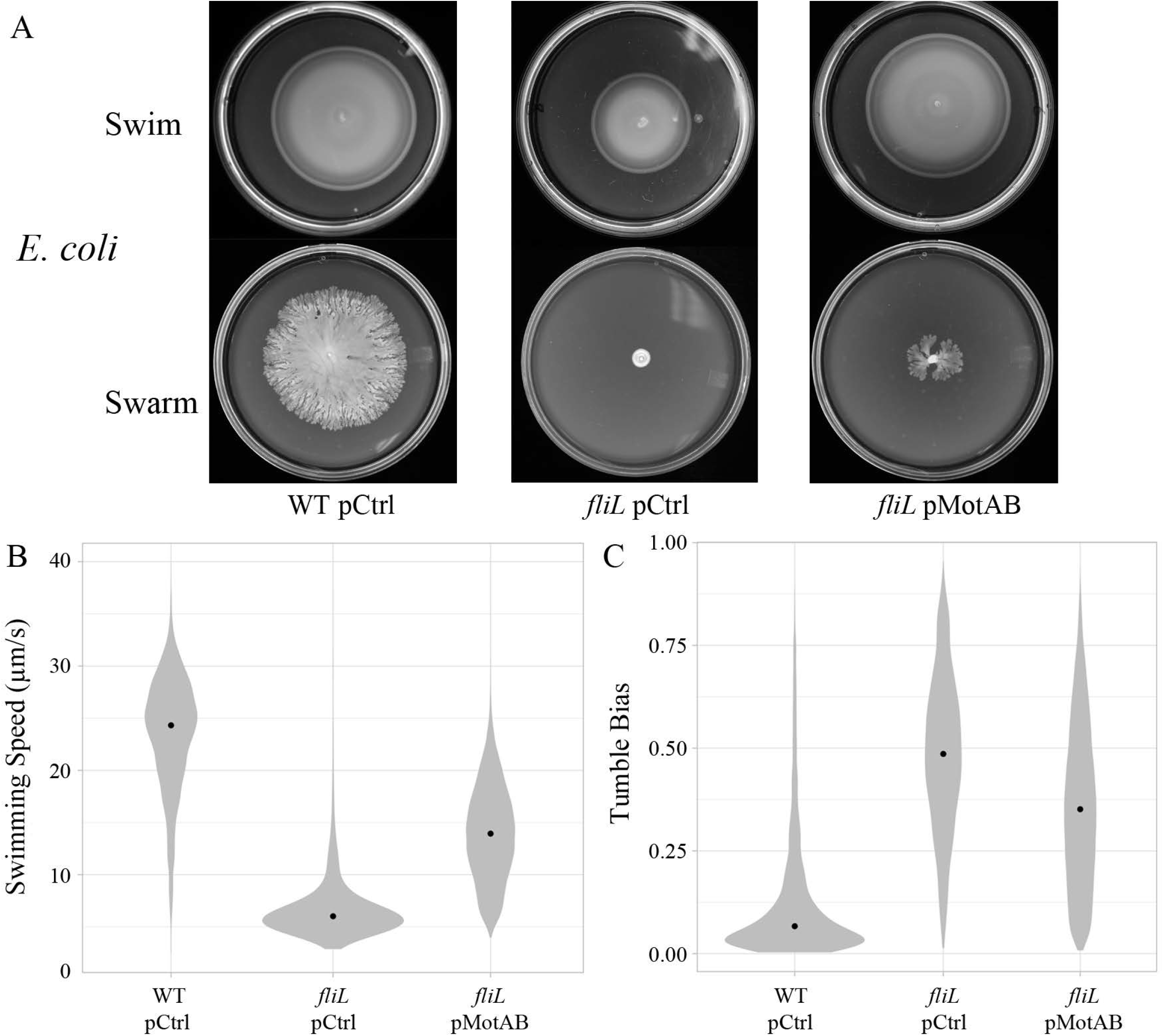
Effects of interspecies FliL exchange on *E. coli* motility. A) Alignment of FliL proteins from *E. coli* (*Ec*), *B. subtilis* (*Bs*), and *P. mirabilis* (*Pm*). Consensus residue matches are colored dark purple, those with a positive BLOSUM62 score are colored pale purple, gaps are white. The N-terminal cytoplasmic domain, predicted trans-membrane (TM) domains, and C-terminal periplasmic domain are indicated. B) *E. coli* WT, and *fliL* strains with either empty plasmid vector (pCtrl), or expressing FliL protein of *E. coli* (FliL^Ec^), *Bacillus* (FliL^Bc^), and *Proteus* (FliL^Pr^) were checked for motility in swim (0.3% agar) and swarm (0.5% Eiken agar) plates supplemented with arabinose. The strains were inoculated in the center and expansion of bacteria (diameter in mm) was recorded after 6 h (swim) or 8 h (swarm), at 30 °C. Error bars are standard deviations from the means. Probability density distributions of C) swimming speeds and D) cell tumble biases of indicated cells lifted from swarms. Each distribution was calculated from more than 5,000 individual trajectories (>1,100 min of cumulative time) for each condition, combined from at least three independent experiments for each strain/condition. Black dot and bars indicate the median and 95% credible intervals of the posterior probabilities of the medians for each treatment. Calculated *p* values are indicated as follows: *, <0.05; **, <0.01; ***, <0.0001; and +, >0.05.

To corroborate results from plate-based complementation of swarming in *E. coli* by FliL from the two other bacteria species, we tracked cells taken from the swarms as before. WT *E. coli* speeds (25.4 µm/s) dropped to 18.5 µm/s in the absence of FliL. pFliL^Ec^ restored speeds to 23.8 µm/s, while pFliL^Bc^ and FliL^Pr^ conferred faster speeds (25.7 and 25 µm/s) (Fig. 3C). WT TB (0.07) increased to 0.27 in the absence of FliL. Complementation with pFliL^Ec^ and pFliL^Bc^ reduced TB to 0.09 and 0.1, respectively, and FliL^Pr^ lowered TB to 0.05 (Fig. 3D). These data appear to suggest that a lower TB is more critical than increased speed for swarm expansion. For example, the speeds of all complemented strains are similar, but the TBs are not; FliL^Pr^, which has the lowest TB, showed the best complementation of *E. coli fliL*.

The interchangeable nature of FliL function during swarming suggests that this function is conserved across phyla.

### Overexpression of stators partially compensates for absence of FliL

We showed earlier that WT *Salmonella* will swarm on higher agar concentrations only when plasmids expressing both FliL and MotAB were provided together (Partridge and Harshey, 2013a). Given the connection of FliL to stator function, we wondered if increased stator levels alone would compensate for the swarming defect of *E. coli fliL*. Overexpression of MotAB proteins from a plasmid only partially restored swarming in the *fliL* mutant (Fig. 4A). We note that in a similar experiment performed with the partially swarming-defective *B. subtilis fliL*, overexpression of MotAB stators also did not compensate the defect (Hall et al., 2018). Cells from the swarm were tracked as before. Compared to the *fliL* mutant speed (17.3 µm/s), MotAB overexpression increased speed (18.4 µm/s), but remained short of WT (25.4 µm/s) (Fig. 4B). The TB of the *fliL* mutant (0.32) decreased with overexpression of MotAB (0.25), but remained well above that of WT (0.07) (Fig. 4B). While it could be argued that FliL overexpression also did not fully restore swarming in the *fliL* mutant, both speeds and TB were nonetheless substantially restored (Fig. 3), effects not seen with MotAB overexpression.

**Figure 4:**
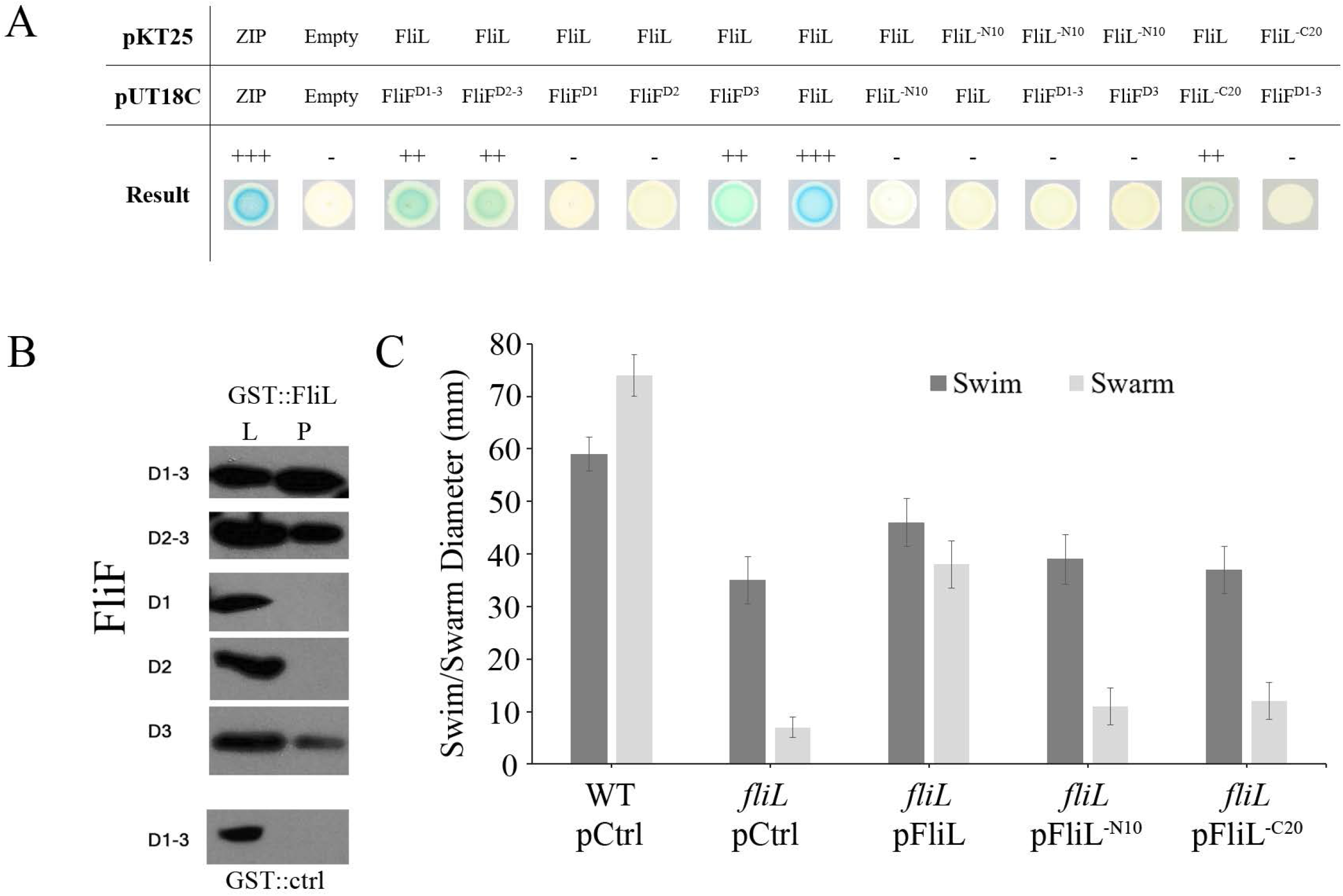
Impact of Stator Overexpression on the movement *E. coli* WT and *fliL*. A) Motility assays of WT *E. coli*, *fliL* pCtrl (empty vector controls) and *fliL* pMotAB (stator overexpression) on swarm plates. Plates were incubated at 30°C for 8 h. Plate assays shown are representative of at least three biological replicates. Probability density distributions of B) swimming speeds and C) cell tumble biases. *E. coli* WT, and *fliL* strains, expressing empty plasmid vector (pCtrl), or expressing MotAB (pMotAB) were grown on LB swarm agar before transfer to LB liquid for observation. Each distribution was calculated from more than 11,000 individual trajectories (>2,300 min of cumulative time) for each condition, combined from five independent experiments for each strain/condition. Black dot and bars indicate the median and 95% credible intervals of the posterior probabilities of the medians for each treatment. Calculated *p* values are indicated as follows: *, <0.05; **, <0.01; ***, <0.0001; and +, >0.05.

We conclude that FliL contribution to motor remodeling is distinct from that of the stators.

### FliL interacts with RBM-3 domain of the MS-ring protein FliF

In *Salmonella*, both pull-down and 2-hybrid experiments had revealed strong FliL-FliL, FliL-MotA, FliL-MotB and FliL-FliF interactions, and comparatively weaker FliL interaction with FliG (Partridge et al., 2015). Here we re-investigated the interaction of FliL with the MS ring protein FliF in *E. coli*. At the time we did this work, bioinformatics analysis of the 560-residue FliF sequence from various bacterial species had identified 3 globular domains in the region spanning residues 43-443 of FliF, referred to as RBM1-3 (Bergeron, 2016). We therefore cloned various combinations of these domains (D), which include the following residues: D1 (43-122), D2 (121-219), and D3 (250-443). The recent cryo-ET structure of the MS ring shows that the N-terminal RBM1 has a TM region, which continues into the periplasm to make up the outer portion of bottom M ring housing an inward facing RBM2 ring, while the bulk of the upper or S ring is RBM3 (Johnson et al., 2020, Tan et al., 2021). From within RBM3 emerges a β-collar that surrounds the rod. RBM3 then loops back into the cytoplasm to interweave with FliG at the top of the C ring switch complex (Lynch et al., 2017, Xue et al., 2018). Thus, RBM3 has a dominant presence in the periplasm, and a smaller, albeit significant one in the cytoplasm. We tested the interaction of these FliF domains with FliL in both 2-hybrid and pull-down assays.

In the BACTH two-hybrid system (Karimova et al., 1998), proteins of interest are genetically fused in various combinations to two fragments (T25 and T18) of the catalytic domain of *Bordetella pertussis* adenylate cyclase (AC) and co-expressed in an *E. coli* strain deficient in endogenous AC. Interaction of the two hybrid proteins brings the T25 and T18 fragments together in the cytoplasm, leading to cyclic AMP (cAMP) synthesis and in turn to transcriptional activation of a *lacZ* reporter system. FliL showed interaction with FliF^D1-3^, FliF^D2-3^, and FliF^D3^, but not FliF^D1^ or FliF^D2^ (Fig. 5A), suggesting FliL interacts only with D3. We confirmed these results by pulldown assays, carried out as previously described (Partridge et al., 2015). Here, N-terminal glutathione S-transferase (GST) was fused to FliL, with various FliF domains co-expressed from inducible plasmids in a *flhDC* strain, ensuring that no other flagellar proteins were present in order to avoid competitive binding to non-tagged proteins. Again, only FliF D3 constructs were reproducibly pulled down with the FliL-GST fusion, as determined by Western blotting (Fig. 5B).

**Figure 5:**
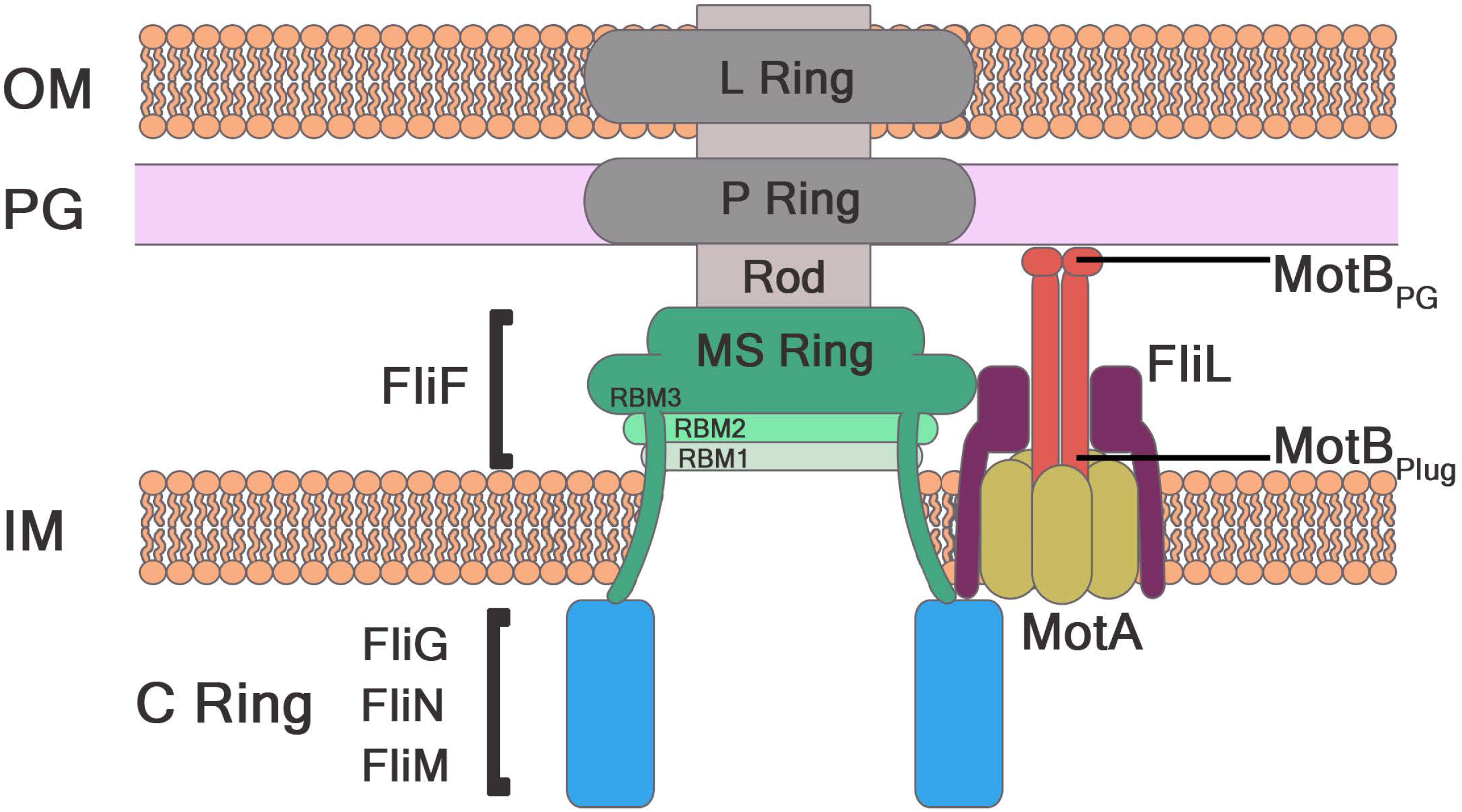
FliL interaction with FliF using two assays. A) Two-hybrid assays using the BACTH system, employing full-length proteins (see Materials and Methods). ZIP, leucine zipper positive control. Negative controls, empty vectors (None). The strength of interactions between the indicated pairs of proteins was measured qualitatively: +++ strong, ++ good, + weak, – none. Images are representative of three independent assays. B) Pulldown assays. A GST fusion of FliL was co-expressed with FliF^D1-3^, FliF^D2-3^, FliF^D1^ or FliF^D2^, and FliF^D3^ in a Δ*flhDC* mutant strain (RP3098). The negative control expressed GST alone. See Materials and Methods for assay details. Lysate controls (L) show protein levels in samples prior to pulldown (P). Assays are representative of at least three independent experiments. C) Motility assays of *E. coli* WT, and *fliL* strains, expressing empty vector (pCtrl), or plasmids expressing *E. coli* FliL or versions missing 10 residues at the N-terminus (FliL^-N10^) or C-terminus (FliL^-C20^). Plates were incubated for 8 h at 30 °C. Error bars are standard deviations from the means.

The N-terminus of *E. coli* FliL is expected to be in the cytoplasm (∼17 residues) with the remaining protein in the periplasm (Attmannspacher et al., 2008) (Fig. 3A). Both the cytoplasmic and periplasmic regions can therefore be expected to contact FliF D3. We generated two truncated versions of *E. coli* FliL, one missing 10 residues at the N-terminus (FliL^-N10^) and the other missing 20 residues at the C-terminus (FliL^-C20^). Using the two-hybrid system, we found that the strong FliL-FliL interaction seen with WT protein was eliminated when one of the partners was FliL^-N10^. Absence of the N-terminus also abrogated interaction with both FliF^D1-3^ or FliF^D3^ (Fig. 5A). These data suggest that the FliL N-terminus interacts with the MS ring. When the C-terminal 20 residues were missing, FliL^-C20^ showed a slightly weaker interaction with full-length FliL compared to FliL-FliL interaction. FliL^-C20^ was also unable to interact with FliF^D1-3^. The 20 C-terminal residues form the last β3 strand of FliL, which is one of the most conserved regions in the protein (Tachiyama et al., 2022, Takekawa et al., 2019, Guo et al., 2022). The data suggest that this region is important for interaction with FliF^D3^ in the periplasm, although we cannot rule out that the C-20 deletion altered the fold of the protein.

Finally, we complemented a Δ*fliL* strain for both swim and swarm motility with the truncated FliLs. While the full-length protein restored motility to the expected suboptimal level (see also Fig. S4B), neither of the truncated versions did (Fig. 5C). FliL^-N10^ is not expected to insert in the membrane. Assuming that FliL^-C20^ folds and inserts correctly in the membrane, the motility data reveal the importance of the C-terminal β3 strand for FliL function, consistent with loss of motor remodeling in the FliL^Pr^ variant FliLc (Fig. 2).

In summary, the strong interaction of FliL with FliF ^D3^, and loss of this interaction in both FliL^-N10^ and FliL^-C20^, suggests that FliL likely interacts with the D3 segment of the MS ring, which is located both in the periplasm and the cytoplasm.

## DISCUSSION

This study reports three new findings with regard to FliL. First, FliL is required for motor remodeling in *E. coli*, *B. subtilis* and *P. mirabilis*, demonstrating a conserved swarming function across phyla. Second, FliL^Ec^ function can be substituted efficiently by FliL^Pm^, and modestly by the phylogenetically distant ortholog FliL^Bs^, showing that FliL performs an equivalent function in all three bacteria. Third, FliL^Ec^ interacts strongly with the RBM-3 domain of the MS ring protein FliF. We discuss these results in the context of what is known about FliL function in bacteria.

### FliL positioning and motor remodeling

<u>FliL positioning at the *E. coli* motor</u>. *fliL* is typically either the first gene in flagellar operons that also transcribe the C ring genes *fliM* and *fliN* or is adjacent to stator genes *motA* and *motB* within the same operon, suggesting that FliL likely interacts with key components of torque– and rotor-bias generating complexes. Based on available structures and positioning of the periplasmic FliL ring around MotAB stators in three different bacteria (Takekawa et al., 2019, Guo et al., 2022, Tachiyama et al., 2022), and on data from this study showing a strong interaction of *E. coli* FliL with RBM-3 of FliF (Fig. 5), we first present a model for FliL positioning at the *E. coli* motor in order to facilitate discussion of the various FliL phenotypes reported in this and in other studies (Fig. 6). In the model, FliL interacts with FliF RBM-3 in both the periplasm and the cytoplasm. The periplasmic interaction with RBM-3 is proposed because the bulk of RBM-3 resides there, and because earlier pull-down assays with only the periplasmic portion of FliF^Se^ showed a strong interaction with FliL^Se^ (Partridge et al., 2015). In the current study, loss of the N-terminal cytoplasmic residues of *E. coli* FliL (FliL^-N10^) results in loss of interaction with RBM-3 in the two-hybrid assay (Fig. 5), suggesting that the FliL N-terminus may interact with RBM-3 C-terminus in the cytoplasm. The combined pull-down and two-hybrid data implicate FliL interaction with FliF in both the periplasm and cytoplasm. Support for interaction of RBM-3 and FliL in the cytoplasm also comes from studies in *Salmonella*, where a weak interaction of FliL with cytoplasmic FliG was seen (Partridge et al., 2015); FliG interdigitates with RBM-3 (Johnson et al., 2020, Bergeron, 2016). The proposed cytoplasmic location of FliL would also place the FliL N-terminus in proximity to MotA residues in the cytoplasm, whose contact with FliG controls not only torque, but also bi-directional rotor switching (Guo and Liu, 2022).

**Figure 6:**
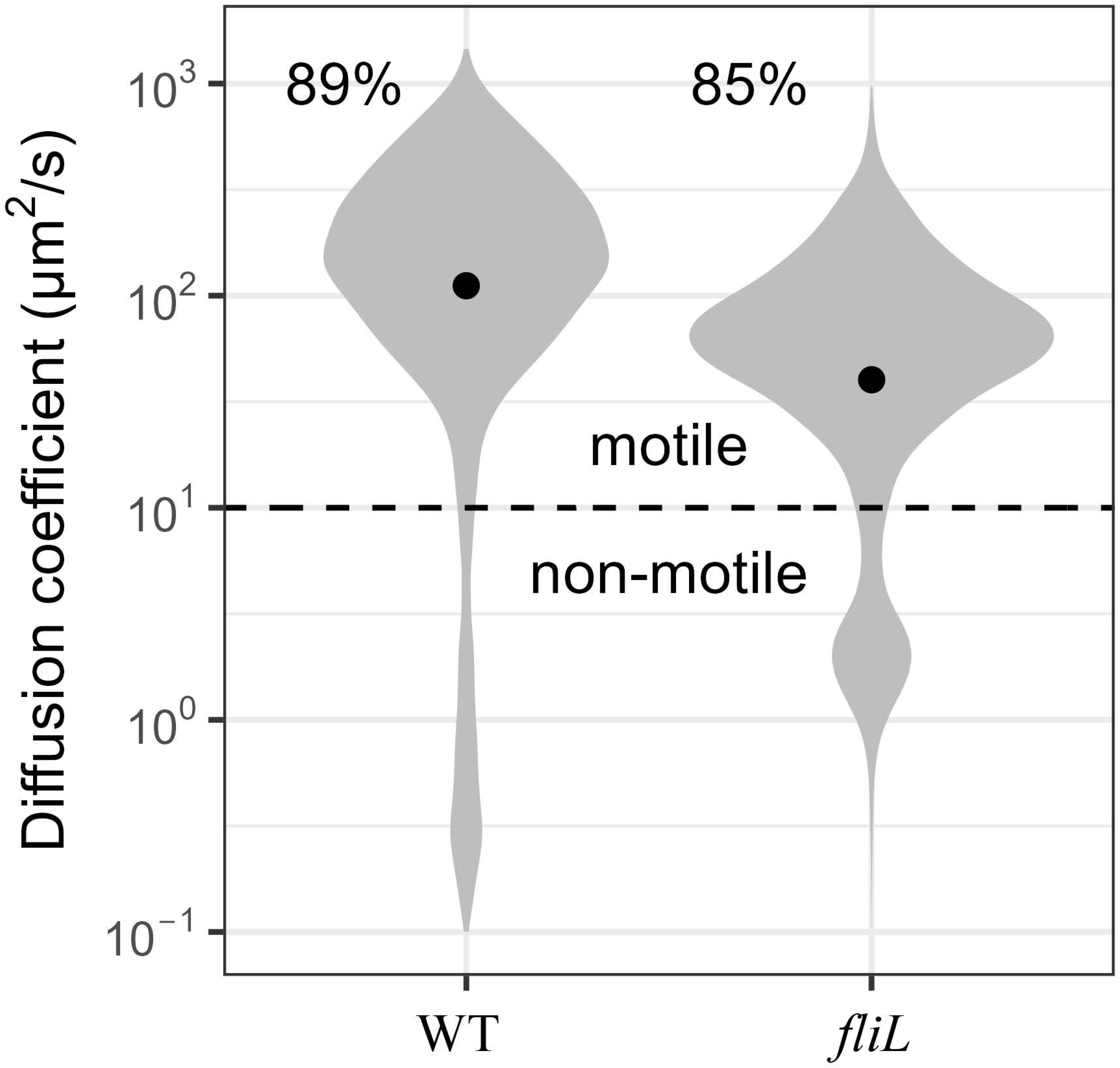
A Model for FliL positioning at the motor in *E. coli/Salmonella*. Based on the data in this and prior studies (see text), we propose that in the periplasm, the multimeric FliL ring interacts with the RBM-3 ring, which interdigitates with the proximal rod above and with FliG in the C-ring below. Loss of this interaction in both FliL^-N10^ and FliL^-C20^, suggests that FliL likely interacts with D3 residues in both periplasm and cytoplasm. A weak interaction of FliL with FliG was reported previously (Partridge et al., 2015).

The model proposed in Fig. 6 is supported by data from other bacterial species. In *B. burdorgofori*, the FliF ring is surrounded by a collar not found in *E. coli*; the FliL ring rests against this collar while encircling the periplasmic ‘plug’ segment of MotB2 beneath its PGB-domain (Peptidoglycan binding) and above MotA5 in the inner membrane (Guo et al., 2022). The cryo-ET images of *B. burdorgofori* basal bodies did not have enough structural resolution to determine if FliL interacts with FliF in the cytoplasm (Guo et al., 2022), but given that the FliL ring is stacked on top MotA5 in the periplasm, it is reasonable to expect the cytoplasmic FliL-N terminus to lie between FliF and MotA. FliL-FliF proximity is also suggested in *C. crescentus*, where developmental degradation of FliF requires FliL (Jenal et al., 1994).

In summary, we propose that in the periplasm, the FliL ring surrounding the stators rests against the RBM-3 ring, which interdigitates with the proximal rod above (Johnson et al., 2020, Tan et al., 2021). In the cytoplasm, FliL interacts with the RBM-3 segment that interdigitates with FliG at the top of the C-ring. FliL likely also contacts FliG, which is in contact with MotA. The deduced location of FliL at the RBM-3-FliG-MotA junction, could allow FliL to influence rotor bias. Thus, FliL is a hub that connects and stabilizes the basal body and stator complexes (Fig. 6).

<u>Motor Remodeling</u>. *E. coli* remodels its motor during swarming (Partridge et al., 2019). When lifted out of the swarm and transferred to liquid, the remodeled motor behavior persists for at least a generation. Motor remodeling was also seen in five other bacterial species – *P. mirabilis, Serratia marcescens, S. enterica, Bacillus subtilis*, and *P. aeruginosa* (Partridge et al., 2020). In the present study, tracking data from cells picked from an *E. coli* swarm, as well as single motor behavior of these cells showed loss of motor remodeling in the absence of FliL (Fig. 1). A similar loss of motor remodeling was observed for the *fliL* null mutant of *B. subtilis,* as well as the *flilc* variant of *P. mirabilis* (Fig. 2), both of which showed suboptimal swarming compared to WT (Fig. S3). Despite the different swarm phenotypes of *fliL* mutants in the three bacteria examined, cell tracking experiments showed that FliL contributed to motor remodeling in all three, and hence had a similar effect at the motor. FliL not only increases motor torque, but also impacts rotor bias, the latter likely to have a large impact on swarming because tumble suppression is expected to improve swarming by fostering side-by-side cell arrangement and pack-like behavior.

The finding that increased expression of MotAB reduces TB, albeit partially, indicates a stator contribution to TB as well (Fig. 4). This is not surprising in the face of recent results showing that the stators and the rotor function as interlocked gears (Santiveri et al., 2020, Deme et al., 2020, Tan et al., 2021). Genetics has also shown that non-functional mutations in rotor proteins are compensated for by suppressors in stator proteins and vice versa (Togashi et al., 1997, Garza et al., 1996, Terashima et al., 2022, Kojima et al., 2011). An important take-home message from this section is that the C ring is not the only player determining rotor bias; the stator complexes, which now include FliL, contribute as well.

### A common function of FliL across phyla

FliL not only plays a similar motor remodeling role in swarming of all three bacterial species studied here, but this function is interchangeable between these species (Fig. 3B-D). *E. coli* and *B. subtilis* are phylogenetically distant (Kunisawa, 1995). *B. subtilis* is a Gram-positive bacterium, whose flagellar basal structure is housed within a cell wall instead of in the periplasm (Mukherjee and Kearns, 2014), and to our knowledge there are no reports of functional complementation of flagellar proteins between the two bacteria. In *B. subtilis*, *fliL* is located immediately upstream of *motAB*, but overexpression of MotAB did not rescue the *fliL* defect (Hall et al., 2018). Instead, a gene *swrD* immediately upstream of *fliL* was essential for swarming; the *swrD* defect could be rescued by overexpression of MotAB. The authors inferred that FliL and SwrD both increase flagellar power, but may do so by different mechanisms. We note that MotAB expression also did not rescue the *fliL* defect in *E. coli* (Fig. 4).

In *P. mirabilis*, where swarmer cells have a distinct hyper-elongated and hyper-flagellated morphology, a Δ*fliL* mutant could swarm, but showed a temperature-sensitive and agar hardness-sensitive swarming behavior (Lee and Belas, 2015). The *P. mirabilis fliL* mutant used in the present study was not a null, but rather had non-WT sequences in the conserved C-terminal β3 strand of FliL (*fliL*c) (Lee and Belas, 2015). While cell tracking experiments showed that FliL contributed to motor remodeling in both *Bacillus* and *Proteus* species (Fig. 2), the cross-species complementation of the *E. coli* Δ*fliL* mutant for swimming and swarming by both FliL^Pm^ and FliL^Bs^, with near WT levels of rescue by FliL^Pm^ was unexpected (Fig. 3B and Fig. S4B). The functional complementation observed by swarming assays was reflected in the cell tracking assays (Fig. 3C-D). Robust complementation for swarming with FliL^Pm^ was accompanied by TB values even lower than those of WT *E. coli*, suggesting that a lower TB may be more critical than speed for swarm expansion (Fig. 3C-D). The similar motor properties displayed across phyla cement the motor remodeling function of FliL.

### FliL contribution to flagella structure and surface sensing

<u>Contribution to flagella structure</u>. In addition to increasing motor torque, FliL contributes to flagella structure/orientation in four different bacterial species – *E. coli, Salmonella*, *C. crescentus*, *B. burgdorfori* – only two of which swarm. In both *E. coli* and *Salmonella*, which sport peritrichous flagella, absence of FliL results in fracture of the rod in a substantial cell population only on swarm media; rod fracture requires that the motor be rotation-proficient (Partridge et al., 2015, Attmannspacher et al., 2008). We have suggested that rod fragility results from higher torque transmitted to the rod during swarming, implicating FliL in stabilizing the rod. We note in this regard that the cryo-ET structure of the *Salmonella* basal body shows rod subunits interdigitated with and packed against the inside of the MS ring β-collar (Johnson et al., 2021, Tan et al., 2021). The rod is made up of 4 different proteins, with layers of interactions between the rod and the MS ring. These layers presumably exist to allow some ‘give’ and provide structural stability to a rotating unit. Our placement of FliL at the rod-proximal S ring (Fig. 6), suggests that FliL may help to fortify this structure and increase its stability in the high-load regime of swarming. Single particle cryo-ET images show a non-coaxial relationship between the rod and the MS ring, i.e. the MS is tilted away from the L and P rings that surround the rod (Johnson et al., 2021). This tilt was attributed by the authors to the removal of the constraints imposed by the two membranes, which were absent in the structure. We note that FliL is also absent in this structure.

A connection between FliL and rod stability was also seen in non-swarming *C. crescentus*, which is polarly flagellated. FliL is not only essential for swimming in this bacterium, but also plays an essential role in the swimmer-to-stalk cell transition, where a developmental c-di-GMP signal results in FliL-dependent FliF degradation accompanied by ejection of the filament attached to the hook-proximal portion of the rod (Jenal et al., 1994, Christen et al., 2007). The ends of the ejected filaments look remarkably similar to those of broken filaments in Δ*fliL Salmonella* swarm cells (Attmannspacher et al., 2008).

In non-swarming *B. burdorfori*, there are two sets of multiple periplasmic flagella at both ends of the bacterium. Both the collar and FliL are important for stator assembly and retention (Motaleb et al., 2011, Guo et al., 2022). Absence of FliL not only has a profound impact of stator assembly and motility, but the periplasmic flagella orient abnormally. We speculate that here too, FliL may be required for co-axial alignment of the MS and LP rings via the rod.

<u>Contribution to surface sensing</u>. The periplasmic region of FliL shows remarkable structural similarity to the mammalian stomatin (SPFH) domain which is known to modulate ion transport/channels in eukaryotes (Takekawa et al., 2019, Tachiyama et al., 2022). These type of proteins are linked to the formation of mechanosensors in *Caenorhabditis elegans,* and mechanosensory transduction in mammalian neurons (Goodman et al., 2002, Wetzel et al., 2007), but their specific mechanism is still unknown (Cheng et al., 2018). FliL was proposed to be a surface-sensor in *P. mirabili*s based on inappropriate swarming behavior of a Δ*fliL* mutant on agar surfaces on which the WT did not swarm, implicating a role for FliL in detecting appropriate surface conditions (Lee and Belas, 2015). A decrease in the expression of the flagellar regulon was also observed in the *Proteus flilc* variant used in our study (Cusick et al., 2012).

A role for the polar flagellum as a surface sensor was first proposed in *V. parahaemolyticus*, which synthesizes hundreds of distinctly different lateral flagella that enable swarming on hard agar (McCarter et al., 1988). In *B. diazoefficiens*, which similarly upregulates lateral flagella, the polar flagellum-associated FliL was implicated in upregulation of lateral flagella (Mengucci et al., 2020), similar to FliL-mediated regulation of the flagellar genes in *P. mirabilis* (Cusick et al., 2012). Given that attempts to rotate the flagellum on a swarming surface may be akin to applying an external load on the flagellum, and given that stator complexes in *E. coli* can move in an out of the rotor by sensing external load applied to the flagellum (Tipping et al., 2013, Lele et al., 2013), could FliL be a sensor of external load? Single motor experiments conducted with planktonic cells of WT and *fliL E. coli* showed that recruitment of stators at the basal body when load was applied externally on the filament, did so irrespective of FliL (Chawla et al., 2017) (see also below).

### Some thoughts on conflicting FliL phenotypes reported in *E. coli*

<u>Surface ‘hardness’ properties</u>. In our hands, FliL null mutants of both *E. coli* and *Salmonella* show similar phenotypes – impaired swimming in 0.3% soft agar (’swim’ or ‘chemotaxis’) media, and no swarming on media with 0.5-0.7% agar (’swarm’) plates (Fig. S4) (Attmannspacher et al., 2008). Since FliL affects rotor bias in these bacteria, the reduced motility of *fliL* mutants in soft agar could at least in part be due to sub-optimal chemotaxis. Swarming does not require chemotaxis proficiency in these bacteria (Burkart et al., 1998), but does require that the rotor be able to switch directions (Mariconda et al., 2006). Che^-^ mutants have extreme CW or CCW biases, and will not swarm unless provided with a more hydrated surface, leading to the inference that bidirectional rotation is somehow initially important for hydrating the surface (Wang et al., 2005, Berg, 2005, Partridge and Harshey, 2013a). The Δ*fliL* mutant does not have an extreme rotation bias, so its non-swarming phenotype should be unrelated to that of the Che^-^ mutants. More likely, the mutant is unable to generate requisite motor output, and suffers rod fracture as cells attempt to move as discussed above. However, a wet surface is also favored by *fliL* mutants, because spraying the swarm agar surface with water provides partial rescue of swarming in the *E. coli* mutant (Fig. S2), and use of the more permission Eiken agar partially rescues swarming in the *Salmonella* mutant (Fig. S2B). We speculate that a ‘wetter’ surface likely lowers the resistance to flagella rotation. It is standard practice in the lab to let swarm plates dry overnight prior to inoculation. If used immediately after pouring, it is not unusual to observe partial swarming in *fliL* mutants. This was the case in the experiments reported by Chawla et al. (Chawla et al., 2017) as well as Lee & Belas (Lee and Belas, 2015), where *fliL* was not observed to affect swarming. *E. coli* is a fastidious swarmer (Partridge and Harshey, 2013b), and can also show sensitivity to different batches of agar obtained from the same commercial source; this again could be a reflection of how much resistance to movement the agar surface provides. Given the clear role of FliL in assisting stator assembly and function in a variety of bacteria, it is not inconceivable that some hard-to-define property of the agar including hydration, influences the difficulty level of swarming and hence the motor output needed for surface navigation. We therefore advise caution when deducing FliL function by simply surveying the literature as was done in a recent report, which concluded that FliL truncations that leave the N-terminal and TM regions intact, block motility, whereas a complete FliL deletion allows normal motility (Liu et al., 2023). This report inferred that the segment of FliL N-terminus that includes the TM region, negatively regulates motor function. We specifically note here that both *E. coli* and *Salmonella* show similar swim and swarm phenotypes irrespective of whether the FliL truncation retains the TM segment (Attmannspacher et al., 2008), or not (see Fig. 3, S2, S4, Table S1).

<u>Swarmer cell membrane composition</u>. The macroscopic morphology (cell length and flagella numbers) of swarmers in some species is dramatically different from their planktonic counterparts (e.g. *P. mirabilis* and *V. parahaemolytics*), comparatively less so in others (e.g. *B. subtilis*), and not strikingly obvious in some others (e.g. *E. coli/Salmonella/Pseudomonas sp*) (Partridge and Harshey, 2013b). However, in cases of Gram-negative species where RNA and protein profiles have been compared, cell physiologies are distinctly different between swarmer and planktonic states (Wang et al., 2006, Morgenstein et al., 2010, Overhage et al., 2008, Gode-Potratz et al., 2011, Pearson et al., 2010, Bhattacharyya et al., 2020). A common observation in swarmers is up-regulation of genes involved in energy metabolism, and changes in membrane lipids and proteins. Given that flagella are embedded in membranes in Gram negative bacteria, with L and P rings supporting the rod in the LPS and PG regions, respectively (Fig. 6), changes in membrane composition or in periplasmic space would be expected to impact how flagella orient/anchor in swarm cells. Such changes could well be responsible for changing FliL positioning at the rotor, producing a pronounced low TB in swarmers. While there is no direct evidence that FliL detects external signals in *E. coli* or *Salmonella*, it is located at an opportune position to link signals coming down from the external filament to the rod, conveying them to the stators and to the cytoplasmic C-ring (Fig. 6). If FliL does indeed have a surface-sensing function, and if that function involves increasing stator recruitment (two big ‘ifs’), then perhaps a proper test for such a function should be conducted in swarm, not in planktonic cells (Chawla et al., 2017).

## EXPERIMENTAL PROCEDURES

### Strains and growth conditions

All strains and plasmids used in this study are described in Supplementary Table 2.

All strains were cultured in Lennox broth (LB; 10 g/liter tryptone, 5 g/liter yeast extract, 5 g/liter NaCl). Starting from single colonies isolated on agar plates, cells were grown to saturation overnight in broth cultures, sub-cultured using a 1:100 dilution ratio in fresh medium for ∼4 h to reach an optical density of 0.4 at 600 nm (OD^600^), and used to inoculate swim or swarm plates. Swim plates (LB solidified with 0.3% Fisher agar) were poured and used on the day of preparation. For the majority of experiments, LB swarm agar plates were supplemented with 0.5% glucose and solidified with 0.5% Eiken agar [Eiken Chemical Co., Japan], while some experiments with *Salmonella* used Fisher agar without glucose. Swarm plates were poured and held at room temperature for 16 h prior to inoculation. Where required, swarm plated were hydrated (supplemental water spritzed onto the surface after drying) as previously described (Wang et al., 2005). Inducer concentrations were arabinose at 0.5% w/v and IPTG at 50 µM. Antibiotic concentrations were: ampicillin (100mg/ml), chloramphenicol (20mg/ml), and kanamycin (50mg/ml).

### Genetic manipulations

DNA was isolated and manipulated by conventional methods (Sambrook, 1989) with genes amplified from the chromosome of WT bacterial species (see Table S1), using appropriate oligonucleotides (available upon request). The coding region of each gene was engineered into either pBAD30, pBAD33, or pBAD18-Kan (Guzman et al., 1995) and verified by DNA sequencing. Deletion of genes was achieved by the one-step mutagenesis procedure (Datsenko and Wanner, 2000). Mutations were transferred to fresh backgrounds by P1*vir* transduction where necessary and verified by PCR amplification. Mutations were converted to unmarked deletions by using the recombinase system of pCP20 (Datsenko and Wanner, 2000). In *Salmonella,* mutations were moved between strains using P22 transduction.

JP2891, the *P. mirabilis fliLc* mutant, was generated by plating the non-motile Tn5-disrupted *fliL* mutant BB2204 on swarm plates and isolating spontaneous *fliL* Swr+ mutants as previously described (Cusick et al., 2012). JP2891 had a *fliL* genotype identical to that described previously YL1001 (Cusick et al., 2012), i.e. loss of Tn5 left the following 68 bp scar sequence –CTGACTCTTATACACAAGTGCGGCCGCGGCCTAGGCGGCCAGATCTGATCAAGAGACAGAATCTGAAC – which shifted the reading frame and altered the last 28 residues of the C-terminus. This was determined by genome sequencing at the GSAF core facility using the HiSeq 4000 platform (PE 2 × 150 setup). The data were analyzed using the breseq program (Deatherage and Barrick, 2014).

### Time-lapse microscopy, cell tracking, and trajectory analysis

Tracking experiments were carried out as described previously (Partridge et al., 2019, Partridge et al., 2020). A custom MATLAB (Mathworks) code (github.com/dufourya/SwimTracker) was used to reconstruct the cell trajectories, and extract speed and tumble bias for single-cells as described previously (Dufour et al., 2016). The subpixel resolution coordinates of each cell in each frame were detected using a previously described method that uses radial symmetry (Parthasarathy, 2012) with an intensity detection threshold set to a false-discovery rate of 5%. Coordinates were linked to obtain cell trajectories using a previously described self-adaptive particle tracking method, u-track 2.1 (Jaqaman et al., 2008), with motion model linkage cost matrices that combine constant velocity and random reorientation with an expected particle velocity of 20 to 30 μm/s and, otherwise, default parameters. Trajectories shorter than 5 s were discarded. The effects of strains and growth conditions on the median swimming speeds and tumble biases were estimated using Bayesian inference with a linear mixed-effect model using the brms library (Bürkner, 2017). Biological replicates were treated as uncorrelated random factors. A log normal and beta distributions were used to model swimming speed and tumble bias, respectively. Bayesian sampling was done with 8 chains with 1,250 iterations each with default parameters. Statistical significance was calculated empirically from the posterior probability distributions of each parameter. Plots were generated in the R statistical environment using the ggplot (Dufour et al., 2016) and tidybayes (Parthasarathy, 2012) libraries.

### Measurement of single motor rotation

Single-motor rotation experiments were performed, and data analyzed, as described previously (Partridge et al., 2015, Partridge et al., 2019), using *E. coli* cells expressing FliC^st^ from plasmid pFD313 (Kuwajima, 1988). Torque output of motors was calculated as described previously (Ryu et al., 2000).

### Two-hybrid screen of protein-protein interactions

Interactions between proteins of interest were screened using the BACTH bacterial two-hybrid system (Euromedex) as described by Karimova et al., (Karimova et al., 1998) and used previously (Partridge et al., 2015). The pKT25 and pUT18C plasmids were used with the host strain BTH101, with a deletion of the flagellar master regulon gene *flhC* (JP319) to prevent native expression of flagellar genes and assembly of flagellar structures. Positive controls utilized the plasmids pKT25-ZIP and pUT18C-ZIP, with each expressing fusion proteins that associate through dimerization of leucine zipper motifs expressed. Negative controls were empty vectors. Genes of interest were amplified from *E. coli* genomic DNA by PCR and fused in frame to the gene fragments in plasmids pKT25 and/or pUT18C. Plasmid pairs were co-transformed into the host strain and plated on LB agar supplemented with ampicillin, kanamycin, X-Gal (5-bromo-4-chloro-3-indolyl-β-D-galactopyranoside; 40 µg/ml^−1^), and IPTG (isopropyl-β-D-thio-galactopyranoside; 0.5 mM). Plates were incubated at 30°C for 48 h, with non-interacting proteins remaining white and potential interactants exhibiting a range of blue coloration.

### GST pulldown assays

Protein pulldown assays were carried out as previously described (Partridge et al., 2015, Paul et al., 2010). Protein pairs to be tested were co-expressed in an *flhDC* strain (RP3098) so that the two proteins of interest were the only flagellar proteins present. Plasmid pHT100 (Tang et al., 1996) was used to express the GST::FliL fusion, with each of the various FliF proteins co-expressed from pTRC99a (NovoPro Bioscience). Control experiments used empty pHT100 (GST only). Immunoblotting used monoclonal anti-FliF antibodies (a gift from May Macnab) and HRP-conjugated anti-mouse antibodies (Sigma).

### Protein Sequence Analysis

Sequence alignments (Fig. 3A) were performed using T-Coffee (Notredame et al., 2000) and edited in Jalview (Waterhouse et al., 2009). The cytoplasmic/transmembrane/periplasmic regions were predicted using DeepTMHMM (Hallgren et al., 2022).

### Supplementary Videos

Referenced recordings of WT and *fliL* cell population movement, transferred from swarm surfaces to liquid, can be found at https://github.com/jpartridge1701/WT-FliL-Swr-Liq.

## ACKNOWLEDGMENTS

We thank graduate students Nabin Bhattaraj and Brady Wilkins for independently confirming the non-swarming phenotype and the genotype of *E. coli fliL* mutants. We thank Daniel Kearns, Philip Rather, and Kelly Hughes for providing strains. This work was supported by Public Health Service Grant GM118085 to R.M.H.

## AUTHOR CONTRIBUTIONS

J.D.P. and R.M.H. conceptualized the study. J.D.P. and R.M.H. designed experiments, J.D.P., R.M.H., and Y.S.D. analyzed the data, J.D.P., R.M.H., and Y.S.D. wrote the manuscript. J.D.P and Y. H. performed experiments.

## GRAPHICAL ABSTRACT

See Fig. 6.

## ABBREVIATED SUMMARY

The flagellum is implicated in surface-sensing since the discovery of *Vibrio parahaemolyticus* swarming decades ago. FliL is a conserved flagellar protein that associates with stators and increases motor torque in several bacteria, and is hypothesized to sense the surface in *P. mirabilis* and to stabilize the flagellar rod in *E. coli* and *Salmonella*. We report here a conserved function for FliL in altering both motor output and rotation bias during swarming. The strong interaction of FliL with the largest domain of the MS ring, which attaches to both the periplasmic rod and cytoplasmic C-ring, places FliL at an opportune position for contributing to rod stability, and for affecting rotor bias through the MS ring, C ring, or stators.

## FIGURE LEGENDS

**Supplementary Figure 1:**
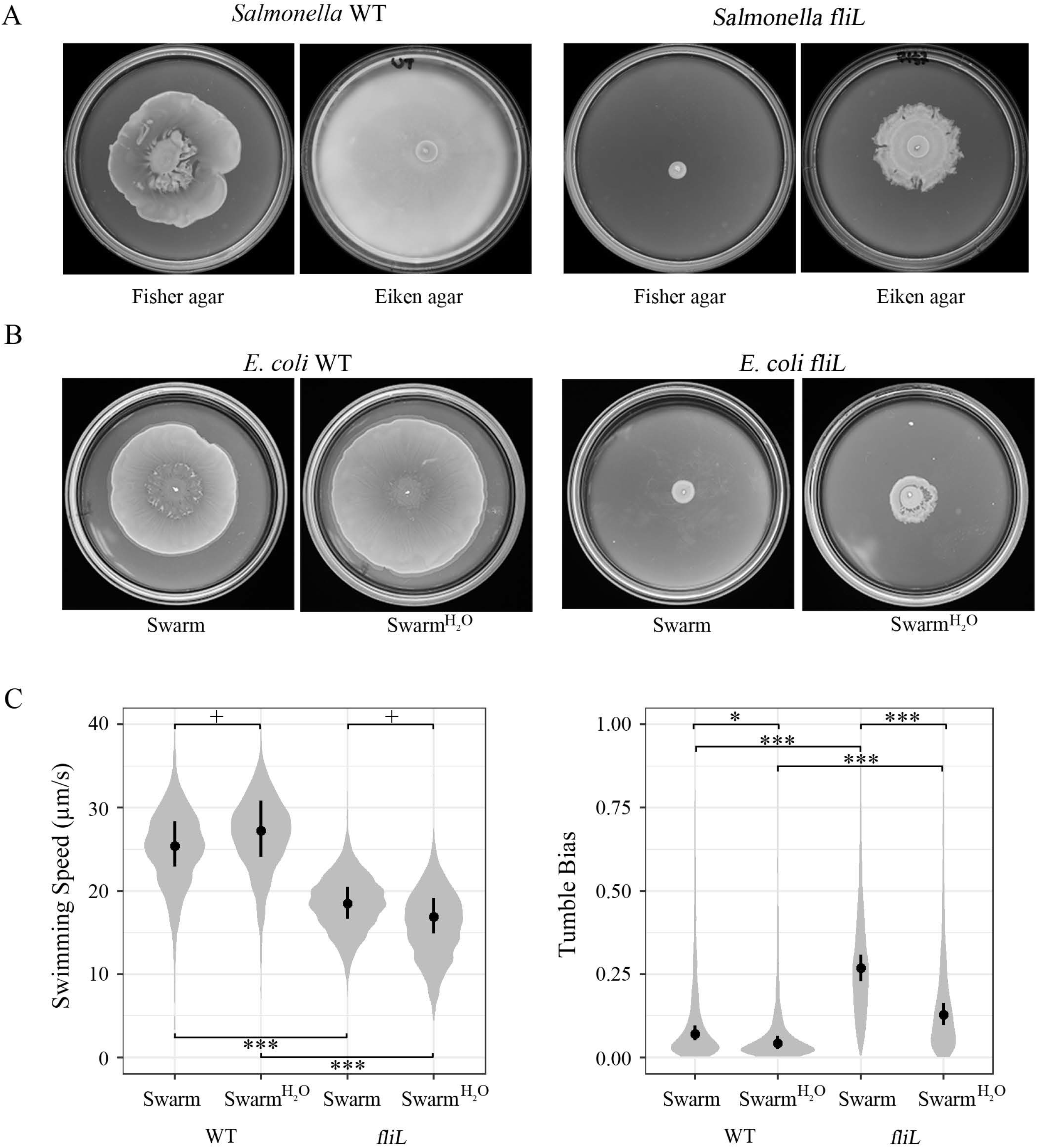
Partitioning of *E. coli* WT and *fliL* swarmers into motile vs. non-motile populations. Shown are the probability density distributions for the cell diffusion coefficients for each strain. Below 10µm^2^/s, cells are labeled as non-motile because they are only subject to Brownian motion. Cells were grown on LB swarm agar before transfer to LB liquid for observation, as described for Fig. 1A. The distribution is representative of a typical tracking experiment for each strain, and is calculated from more than 10,000 individual trajectories (>2,000 min of cumulative time) for each. The diffusion coefficient is reflective of motion models that account for directed motion and random reorientation with an expected particle velocity (see Materials and Methods).

**Supplementary Figure 2:**
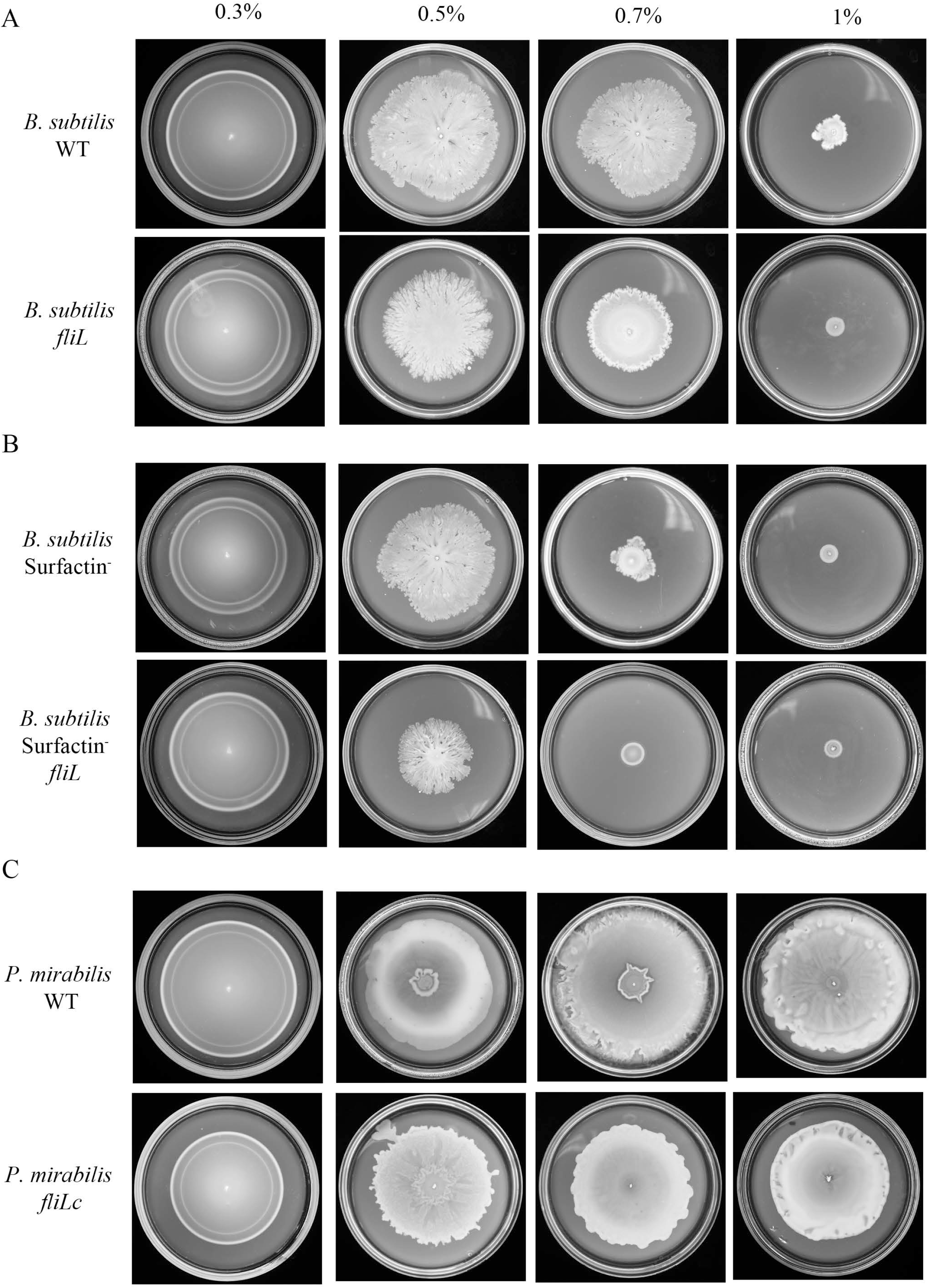
Motility behavior of *E. coli* and *Salmonella* WT and *fliL* strains under more permissive swarming conditions. A) *Salmonella* wild-type (LT2), and *fliL* (TH7357) strains, were inoculated on both Fisher and Eiken agar (0.5% w/v) swarm plates (with 0.5% glucose) and incubated for 12 h at 37 °C. Plate assays shown are representative of at least three biological replicates. B) Swarm assays of wild-type (MG1655) and *fliL* (JP1790) *E. coli* on swarm plates (with 0.5% glucose), with and without extra hydration (see Methods, plates were incubated at room temperature for 15 min after hydration), labelled “Swarm” and “Swarm^+H20^” respectively. C) Probability density distribution of *E. coli* swimming speeds (left) and tumble biases (right) within a pseudo-2D environment. Cells from B were transferred to LB liquid for observation. *fliL* swarms on the ‘H2O’ plates were checked via microscopy to ensure that characteristic swirls associated with collective motion were evident. Each distribution was calculated from more than 7,700 individual trajectories (>1,760 min of cumulative time) for each condition, combined from 3 independent experiments for each. Black dot and bars indicate the median and 95% credible intervals of the posterior probabilities of the medians for each treatment. Calculated *p* values are indicated as follows: *, <0.05; **, <0.01; ***, <0.0001; and +, >0.05.

**Supplementary Figure 3:**
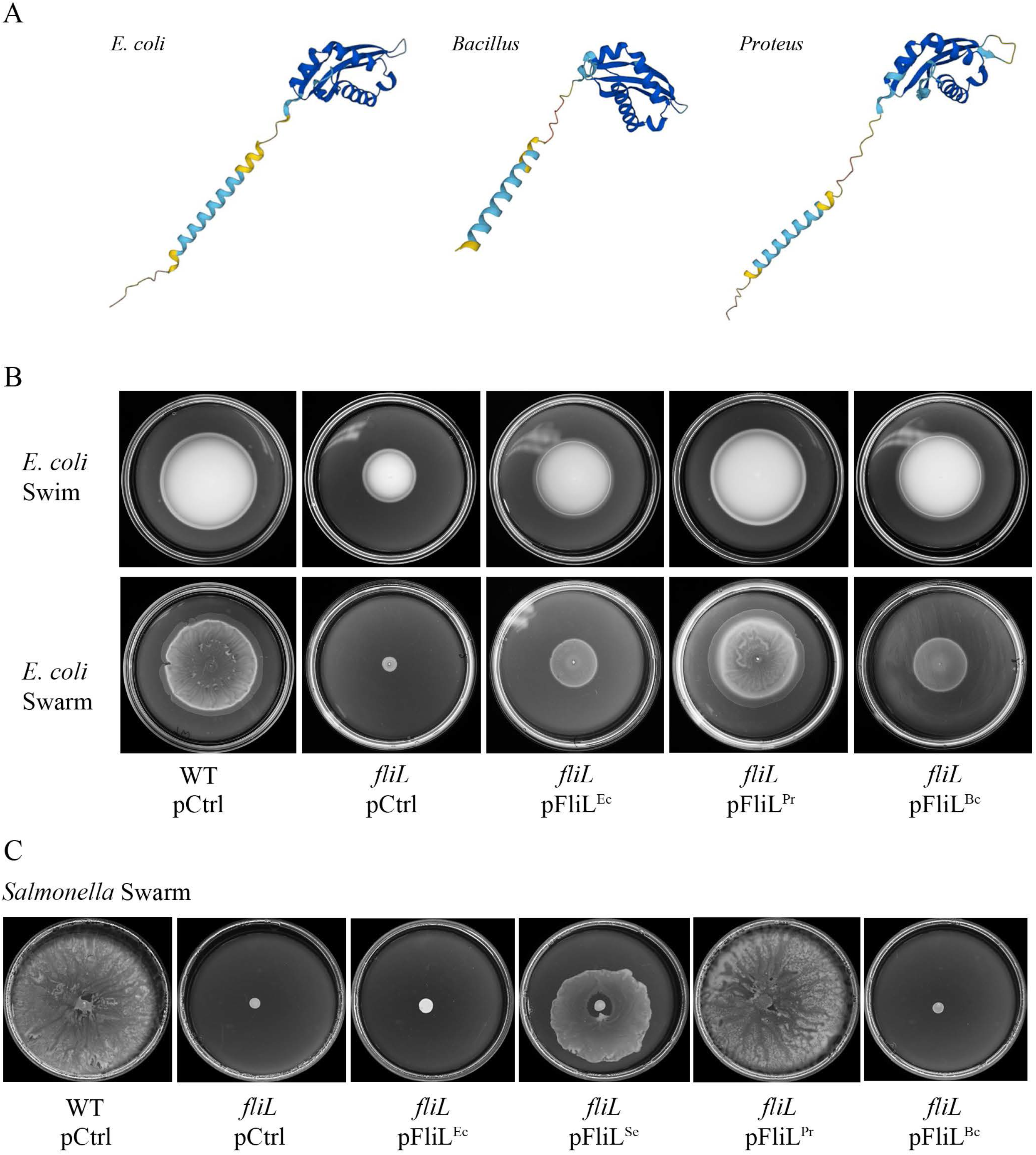
Motility behavior of wild-type and *fliL* strains of *Bacillus subtilis* and *Proteus mirabilis*. Swim (LB solidified with 0.3% agar) and Swarm (LB solidified with 0.5, 0.7, and 1% Eiken agar), assays of wild-type and *fliL* strains of A) *B. subtilis* (3610 and DS6540), B) *B. subtilis* surfactin-strains (DS1122 and DS9132), and C) *P. mirabilis* (BB2000 and JP2891). Cultures were inoculated in the center of each plate and incubated at 30°C for 8 h (swim) or 16 h (swarm). Plate assays shown are representative of at least three biological replicates.

**Supplementary Figure 4:** Predicted protein structures of FliL in *E. coli, B. subtilis, P. mirabilis,* and Effects of interspecies FliL exchange on *E. coli* and *Salmonella* motility. A) Structures predicted and generated using available coding sequence data via KEGG (Kanehisa and Goto, 2000), and the Alphafold predictive algorithm (Jumper et al., 2021, Varadi et al., 2021). B) *E. coli* wild-type, and *fliL* strains, expressing empty vector (pCtrl), or plasmids expressing FliL of *E. coli* (FliL^Ec^), *Bacillus* (FliL^Bc^), and *Proteus* (FliL^Pr^) were checked for motility on swim and swarm plates as described in Fig. 3B. C) *Salmonella* wild-type (14028) and *fliL* (JP1809) strains, expressing empty vector (pCtrl), or plasmids expressing FliL of *E. coli* (FliL^Ec^), *Salmonella* (FliL^Se^), *Bacillus* (FliL^Bc^), and *Proteus* (FliL^Pr^) were inoculated on 0.5% Fisher agar swarm plates supplemented with arabinose, and incubated for 10 h at 37 °C. Plate assays shown are representative of at least three biological replicates.

**Supplementary Table S1:** **Mean posterior probabilities for the median tumble biases and swimming speeds.** Bayesian inference was used to determine if the medians of the swimming speeds and tumble biases are significantly different between liquid and swarm preparations. The posterior probability distributions of the medians for each strain and each treatment were calculated using a linear mixed-effect model with a Gaussian distribution link function. The mean and 95% credible intervals^a^ of the posteriors of the medians for each distribution is also reported. See supplementary information for more details. The means and 95% credible intervals^b^ of the differences of the medians between conditions is reported. *P* values^c^ (for difference in the medians >0 or <0) were calculated by sampling the posterior probability distributions.

|  | Median posteriors<sup>a</sup><br>[95% CI] | Variables | Conditions<br>contrast<sup>b</sup> [95% CI] | *P*<br>value<sup>c</sup> |
| --- | --- | --- | --- | --- |
| **Figure 1A**<br>**Speeds** |  |  |  |  |
| WT Liquid | 22.6 [20.95, 24.44] | Liq vs. Swarm | 0.15 [0.05, 0.26] | 0.004 |
| WT Swarm | 25.38 [23.55, 27.18] | Wt vs. *fliL* | 0.36 [0.25, 0.46] | <0.0001 |
| *fliL* Liquid | 15.85 [14.68, 17.07] | Interaction Strain and<br>Condition | -0.04 [-0.19, 0.11] | 0.29 |
| *fliL* Swarm | 18.51 [17.16, 19.88] |  |  |  |
| **Figure 1A TB** |  |  |  |  |
| WT Liquid | 0.11 [0.08, 0.13] | Liq vs. Swarm | 0.63 [0.34, 0.91] | 0.0004 |
| WT Swarm | 0.07 [0.053, 0.89] | Wt vs. *fliL* | -0.49 [-0.83, -0.16] | 0.0047 |
| *fliL* Liquid | 0.16 [0.13, 0.19] | Interaction Strain and<br>Condition | -1.08 [-1.54, -0.60] | 0.0006 |
| *fliL* Swarm | 0.26 [0.23, 0.30] |  |  |  |
| **Figure 2**<br>***Proteus* Speeds** |  |  |  |  |
| WT Liquid | 7.55 [7.03, 8.17] | Liq vs. Swarm | -0.15 [-0.28, -0.03] | 0.013 |
| WT Swarm | 14.43 [13.46, 15.49] | Wt vs. *fliL* | -0.47 [-0.57, -0.36] | <0.0001 |
| *fliL* Liquid | 12.07 [11.12, 13.01] | Interaction Strain and<br>Condition | 0.81 [0.63, 0.96] | <0.0001 |
| *fliL* Swarm | 10.37 [9.39, 11.42] |  |  |  |
| **Figure 2**<br>***Proteus* TB** |  |  |  |  |
| WT Liquid | 0.23 [0.19, 0.28] | Liq vs. Swarm | 0.15 [-0.24, 0.50] | 0.21 |
| WT Swarm | 0.20 [0.17, 0.24] | Wt vs. *fliL* | -0.43 [-0.76, -0.11] | 0.009 |
| *fliL* Liquid | 0.32 [0.27, 0.37] | Interaction Strain and<br>Condition | -0.30 [-0.78, 0.21] | 0.10 |
| *fliL* Swarm | 0.35, [28, 0.41] |  |  |  |
| **Figure 2**<br>***Bacillus***<br>**Speeds** |  |  |  |  |
| WT Liquid | 18.96 [16.89, 21.30] | Liq vs. Swarm | 0.45 [0.27, 0.65] | 0.0003 |
| WT Swarm | 29.83 [25.64, 34.80] | Wt vs. *fliL* | -0.28 [-0.45, -0.13] | 0.0027 |
| *fliL* Liquid | 14.26 [12.65, 16.21] | Interaction Strain and<br>Condition | -0.23 [-0.47, 0.02] | 0.03 |
| *fliL* Swarm | 17.90 [16.14, 19.66] |  |  |  |
| **Figure 2**<br>***Bacillus* TB** |  |  |  |  |
| WT Liquid | 0.24 [0.20, 0.29] | Liq vs. Swarm | -1.45 [-2.16, -0.93] | <0.0001 |
| WT Swarm | 0.07 [0.035, 0.10] | Wt vs. *fliL* | -0.26 [-0.62, 0.09] | 0.06 |
| *fliL* Liquid | 0.20 [0.16, 0.24] | Interaction Strain and Condition | 1.77 [1.16, 2.56] | <0.0001 |
| *fliL* Swarm | 0.25 [0.22, 0.29] |  |  |  |
| **Figure 3 Speeds** |  |  |  |  |
| WT pCtrl | 25.35 [23.85, 26.90] | WT pCtrl vs. *fliL* pCtrl | 0.32 [0.23, 0.40] | <0.0001 |
| *fliL* pCtrl | 18.46 [17.39, 19.65] | WT Ctrl vs. *fliL* pFliL<sup>Ec</sup> | 0.06 [-0.02, 0.15] | 0.069 |
| *fliL* pFliL<sup>Ec</sup> | 23.75 [22.12, 25.25] | *fliL* pCtrl vs. *fliL* pFliL<sup>Ec</sup> | 0.25 [0.16, 0.34] | 0.0001 |
| *fliL* pFliL<sup>Bc</sup> | 25.72 [24.20, 27.44] | *fliL* pCtrl vs. *fliL* pFliL<sup>Bc</sup> | 0.33 [0.24, 0.42] | <0.0001 |
| *fliL* pFliL<sup>Pr</sup> | 24.99 [22.90, 27.29] | *fliL* pCtrl vs. *fliL* pFliL<sup>Pr</sup> | 0.30 [0.19, 0.41] | 0.0001 |
| **Figure 3 TB** |  |  |  |  |
| WT pCtrl | 0.07 [0.05, 0.09] | WT pCtrl vs. *fliL* pCtrl | -1.57 [-2.00, -1.17] | <0.0001 |
| *fliL* pCtrl | 0.27 [0.23, 0.32] | WT Ctrl vs. *fliL* pFliL<sup>Ec</sup> | -0.35 [-0.84, 0.13] | 0.068 |
| *fliL* pFliL<sup>Ec</sup> | 0.09 [0.07, 0.13] | *fliL* pCtrl vs. *fliL* pFliL<sup>Ec</sup> | -1.22 [-1.63, -0.82] | <0.0001 |
| *fliL* pFliL<sup>Bc</sup> | 0.10 [0.07, 0.13] | *fliL* pCtrl vs. *fliL* pFliL<sup>Bc</sup> | -1.19 [-1.57, -0.81] | <0.0001 |
| *fliL* pFliL<sup>Pr</sup> | 0.05 [0.02, 0.08] | *fliL* pCtrl vs. *fliL* pFliL<sup>Pr</sup> | -2.00 [-2.78, -1.41] | <0.0001 |
| **Figure 4 Speeds** |  |  |  |  |
| WT pCtrl | 25.37 [23.29, 27.47] | WT pCtrl vs. *fliL* MotAB | 0.32 [0.21, 0.43] | <0.0001 |
| *fliL* pCtrl | 17.33 [15.87, 18.80] | WT pCtrl vs. *fliL* pCtrl | 0.38 [0.27, 0.50] | <0.0001 |
| *fliL* pMotAB | 18.41 [17.11, 19.94] | *fliL* pMotAB vs. *fliL* pCtrl | -0.06 [-0.18, 0.05] | 0.13 |
| **Figure 4 TB** |  |  |  |  |
| WT pCtrl | 0.07 [0.04, 0.10] | WT pCtrl vs. *fliL* MotAB | -1.46 [-2.02, -0.93] | <0.0001 |
| *fliL* pCtrl | 0.32 [0.26, 0.38] | WT pCtrl vs. *fliL* pCtrl | -1.81 [-2.38, -1.28] | <0.0001 |
| *fliL* pMotAB | 0.25 [0.19, 0.31] | *fliL* pMotAB vs. *fliL* pCtrl | 0.35 [-0.04, 0.80] | 0.038 |
| **Supplementary Figure 2 Speeds** |  |  |  |  |
| WT Swarm | 25.40 [22.87, 28.20] | WT Swarm vs. WT Swarm<sup>H2O</sup> | 0.07 [-0.10, 0.23] | 0.18 |
| WT Swarm<sup>H2O</sup> | 27.23 [23.88, 30.59] | WT Swarm vs. *fliL* Swarm | -0.32 [0.17, 0.47] | 0.0001 |
| *fliL* Swarm | 18.49 [16.68, 20.51] | *fliL* Swarm vs. *fliL* Swarm<sup>H2O</sup> | -0.09 [-0.26, 0.07] | 0.11 |
| *fliL* Swarm<sup>H2O</sup> | 16.89 [14.78, 19.05] | WT Swarm<sup>H2O</sup> vs. *fliL* Swarm<sup>H2O</sup> | 0.41 [0.25, 0.58] | <0.0001 |
| **Supplementary Figure 2 TB** |  |  |  |  |
| WT Swarm | 0.07 [0.05, 0.09] | WT Swarm vs. WT Swarm<sup>H2O</sup> | -0.55 [-1.17, 0.02] | 0.028 |
| WT Swarm<sup>H2O</sup> | 0.04 [0.02, 0.06] | WT Swarm vs. *fliL* Swarm | -1.57 [-1.96, -1.18] | <0.0001 |
| *fliL* Swarm | 0.27 [0.23, 0.31] | *fliL* Swarm vs. *fliL* Swarm<sup>H2O</sup> | -0.91 [-1.28, -0.56] | <0.0001 |
| *fliL* Swarm<sup>H2O</sup> | 0.13 [0.10, 0.16] | WT Swarm<sup>H2O</sup> vs. *fliL* Swarm<sup>H2O</sup> | -1.21 [-1.83, -0.66] | 0.0002 |

**Table S2:**
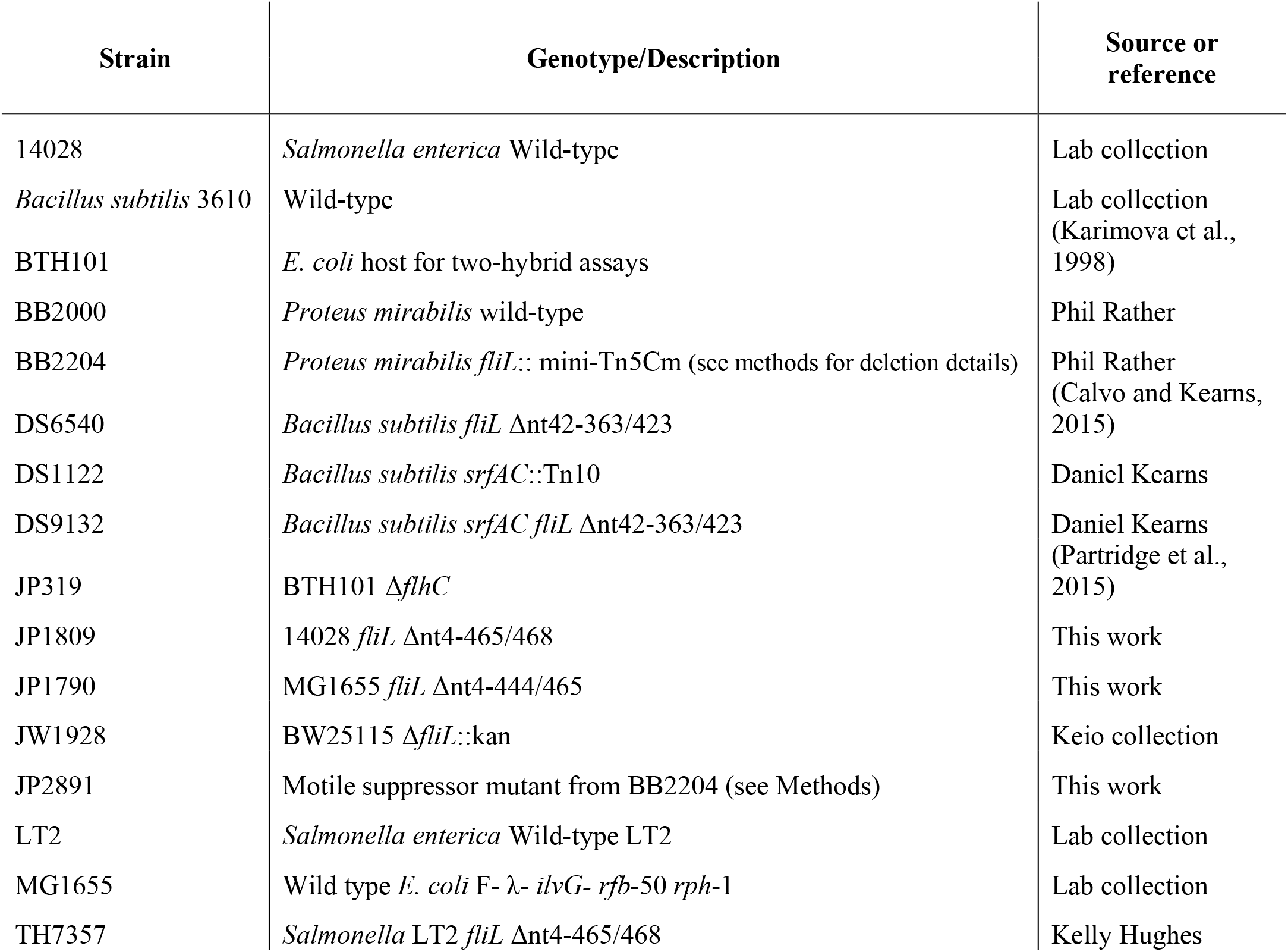
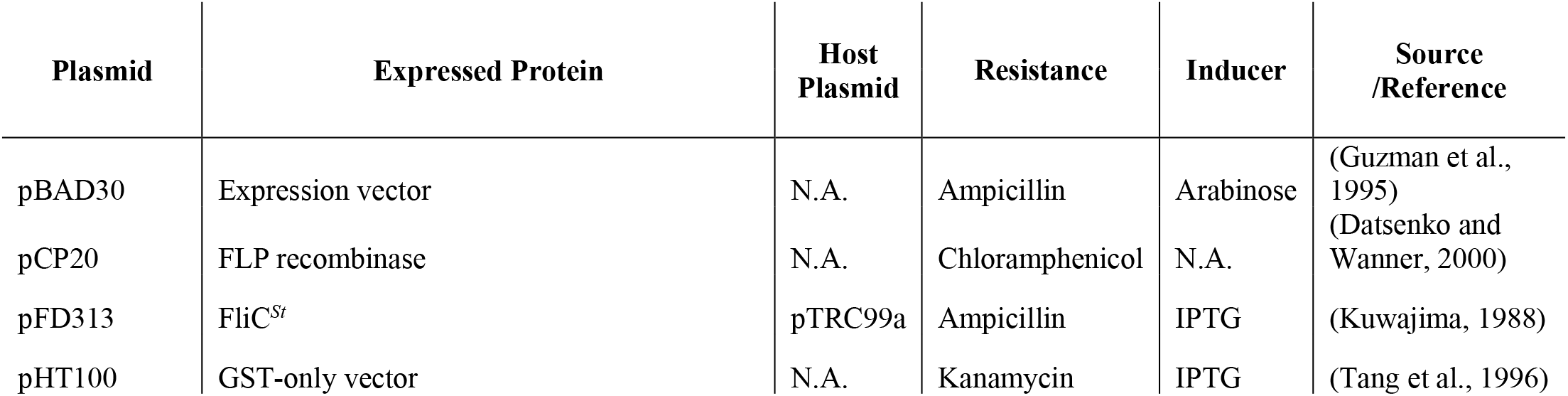
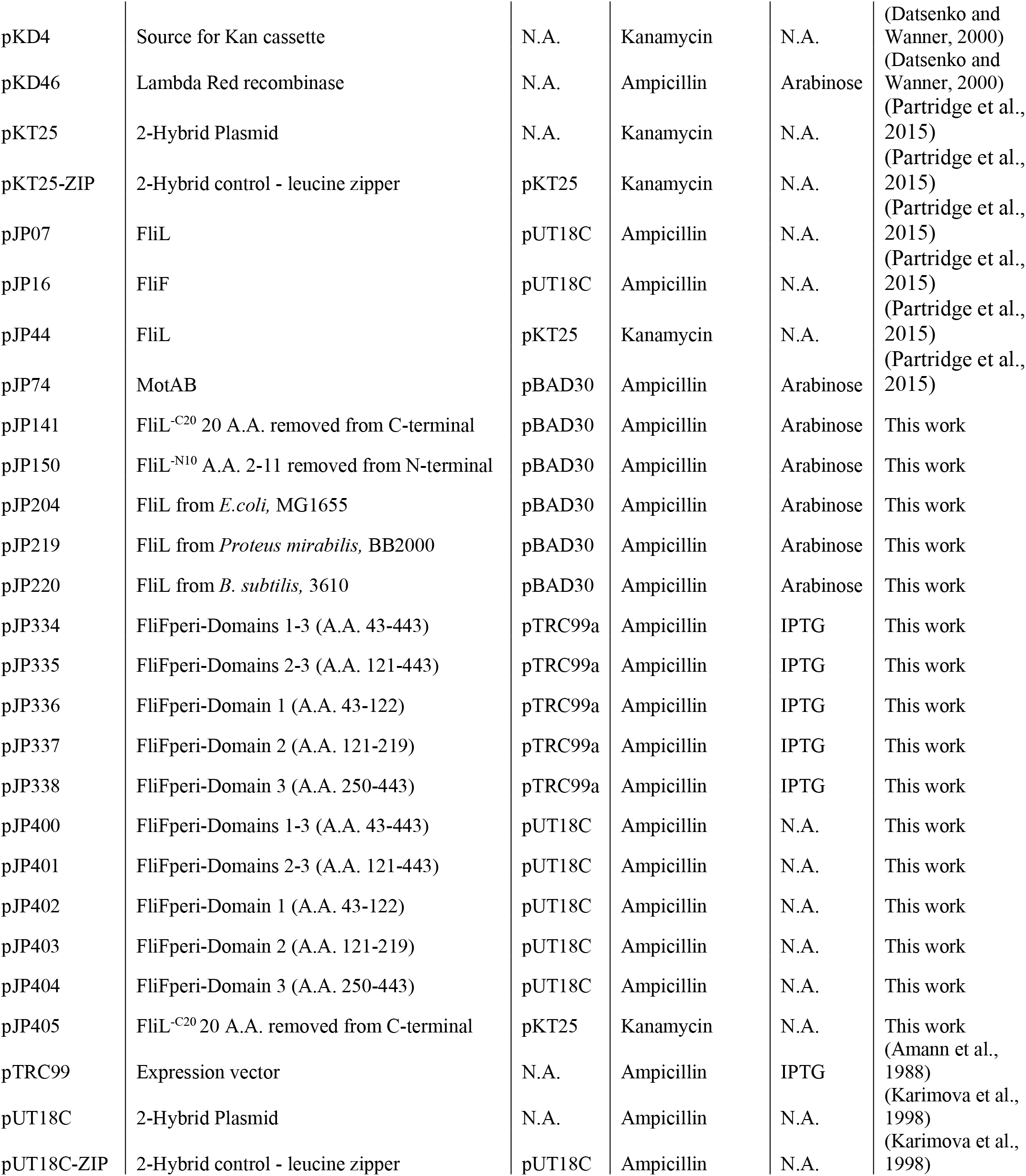
**Strains and Plasmids used in this study.** Under Genotype, the numbers after Δ (deletion), denote the missing nucleotide sequence, and the number after the slash (/) indicates the total sequence length, confirmed by DNA sequencing.

## REFERENCES

1. Attmannspacher, U., Scharf, B. E. & Harshey, R. M. 2008. FliL is essential for swarming: motor rotation in absence of FliL fractures the flagellar rod in swarmer cells of Salmonella enterica. Mol Microbiol, 68, 328–41.

2. Baker, A. E. & O’Toole, G. A. 2017. Bacteria, Rev Your Engines: Stator Dynamics Regulate Flagellar Motility. J Bacteriol, 199.

3. Be’er, A. & Ariel, G. 2019. A statistical physics view of swarming bacteria. Mov Ecol, 7, 9.

4. Berg, H. C. 2003. The rotary motor of bacterial flagella. Annu Rev Biochem, 72, 19–54.

5. Berg, H. C. 2005. Swarming motility: it better be wet. Curr Biol, 15, R599–600.

6. Berg, H. C. & Turner, L. 1993. Torque generated by the flagellar motor of Escherichia coli. Biophys J, 65, 2201–16.

7. Bergeron, J. R. 2016. Structural modeling of the flagellum MS ring protein FliF reveals similarities to the type III secretion system and sporulation complex. PeerJ, 4, e1718.

8. Bhattacharyya, S., Walker, D. M. & Harshey, R. M. 2020. Dead cells release a ‘necrosignal’ that activates antibiotic survival pathways in bacterial swarms. Nat Commun, 11, 4157.

9. Burkart, M., Toguchi, A. & Harshey, R. M. 1998. The chemotaxis system, but not chemotaxis, is essential for swarming motility in Escherichia coli. Proc Natl Acad Sci U S A, 95, 2568–73.

10. Bürkner, P.-C. 2017. brms: An R Package for Bayesian Multilevel Models Using Stan. Journal of Statistical Software, 80, 1–28.

11. Chaban, B., Hughes, H. V. & Beeby, M. 2015. The flagellum in bacterial pathogens: For motility and a whole lot more. Semin Cell Dev Biol, 46, 91–103.

12. Chawla, R., Ford, K. M. & Lele, P. P. 2017. Torque, but not FliL, regulates mechanosensitive flagellar motor-function. Scientific Reports, 7, 5565.

13. Cheng, Y. R., Jiang, B. Y. & Chen, C. C. 2018. Acid-sensing ion channels: dual function proteins for chemo-sensing and mechano-sensing. J Biomed Sci, 25, 46.

14. Chevance, F. F. & Hughes, K. T. 2008. Coordinating assembly of a bacterial macromolecular machine. Nat Rev Microbiol, 6, 455–65.

15. Christen, M., Christen, B., Allan, M. G., Folcher, M., Jenö, P., Grzesiek, S. & Jenal, U. 2007. DgrA is a member of a new family of cyclic diguanosine monophosphate receptors and controls flagellar motor function in Caulobacter crescentus. Proceedings of the National Academy of Sciences, 104, 4112–4117.

16. Cusick, K., Lee, Y. Y., Youchak, B. & Belas, R. 2012. Perturbation of FliL interferes with Proteus mirabilis swarmer cell gene expression and differentiation. J Bacteriol, 194, 437–47.

17. Darnton, N. C., Turner, L., Rojevsky, S. & Berg, H. C. 2010. Dynamics of bacterial swarming. Biophys J, 98, 2082–90.

18. Datsenko, K. A. & Wanner, B. L. 2000. One-step inactivation of chromosomal genes in Escherichia coli K-12 using PCR products. Proceedings of the National Academy of Sciences, 97, 6640–6645.

19. Deatherage, D. E. & Barrick, J. E. 2014. Identification of mutations in laboratory-evolved microbes from next-generation sequencing data using breseq. Methods Mol Biol, 1151, 165–88.

20. Deme, J. C., Johnson, S., Vickery, O., Aron, A., Monkhouse, H., Griffiths, T., James, R. H., Berks, B. C., Coulton, J. W., Stansfeld, P. J. & Lea, S. M. 2020. Structures of the stator complex that drives rotation of the bacterial flagellum. Nat Microbiol, 5, 1553–1564.

21. Duan, Q., Zhou, M., Zhu, L. & Zhu, G. 2013. Flagella and bacterial pathogenicity. Journal of Basic Microbiology, 53, 1–8.

22. Dufour, Y. S., Gillet, S., Frankel, N. W., Weibel, D. B. & Emonet, T. 2016. Direct Correlation between Motile Behavior and Protein Abundance in Single Cells. PLoS Comput Biol, 12, e1005041.

23. Garza, A. G., Biran, R., Wohlschlegel, J. A. & Manson, M. D. 1996. Mutations in motB suppressible by changes in stator or rotor components of the bacterial flagellar motor. J Mol Biol, 258, 270–85.

24. Ghelardi, E., Salvetti, S., Ceragioli, M., Gueye, S. A., Celandroni, F. & Senesi, S. 2012. Contribution of surfactin and SwrA to flagellin expression, swimming, and surface motility in Bacillus subtilis. Appl Environ Microbiol, 78, 6540–4.

25. Gode-Potratz, C. J., Kustusch, R. J., Breheny, P. J., Weiss, D. S. & Mccarter, L. L. 2011. Surface sensing in Vibrio parahaemolyticus triggers a programme of gene expression that promotes colonization and virulence. Mol Microbiol, 79, 240–63.

26. Goodman, M. B., Ernstrom, G. G., Chelur, D. S., O’hagan, R., Yao, C. A. & Chalfie, M. 2002. MEC-2 regulates C. elegans DEG/ENaC channels needed for mechanosensation. Nature, 415, 1039–42.

27. Guo, S. & Liu, J. 2022. The Bacterial Flagellar Motor: Insights Into Torque Generation, Rotational Switching, and Mechanosensing. Front Microbiol, 13, 911114.

28. Guo, S., Xu, H., Chang, Y., Motaleb, M. A. & Liu, J. 2022. FliL ring enhances the function of periplasmic flagella. Proceedings of the National Academy of Sciences, 119, e2117245119.

29. Guzman, L. M., Belin, D., Carson, M. J. & Beckwith, J. 1995. Tight regulation, modulation, and high-level expression by vectors containing the arabinose PBAD promoter. J Bacteriol, 177, 4121–30.

30. Hall, A. N., Subramanian, S., Oshiro, R. T., Canzoneri, A. K. & Kearns, D. B. 2018. SwrD (YlzI) Promotes Swarming in Bacillus subtilis by Increasing Power to Flagellar Motors. J Bacteriol, 200.

31. Hallgren, J., Tsirigos, K. D., Pedersen, M. D., Almagro Armenteros, J. J., Marcatili, P., Nielsen, H., Krogh, A. & Winther, O. 2022. DeepTMHMM predicts alpha and beta transmembrane proteins using deep neural networks. bioRxiv, 2022.04.08.487609.

32. Harshey, R. M. 2003. Bacterial motility on a surface: many ways to a common goal. Annu Rev Microbiol, 57, 249–73.

33. Harshey, R. M. & Matsuyama, T. 1994. Dimorphic transition in Escherichia coli and Salmonella typhimurium: surface-induced differentiation into hyperflagellate swarmer cells. Proc Natl Acad Sci U S A, 91, 8631–5.

34. Homma, M. & Kojima, S. 2022. The Periplasmic Domain of the Ion-Conducting Stator of Bacterial Flagella Regulates Force Generation. Frontiers in Microbiology, 13.

35. Hosking, E. R., Vogt, C., Bakker, E. P. & Manson, M. D. 2006. The Escherichia coli MotAB proton channel unplugged. J Mol Biol, 364, 921–37.

36. Jaqaman, K., Loerke, D., Mettlen, M., Kuwata, H., Grinstein, S., Schmid, S. L. & Danuser, G. 2008. Robust single-particle tracking in live-cell time-lapse sequences. Nat Methods, 5, 695–702.

37. Jarrell, K. F. & Mcbride, M. J. 2008. The surprisingly diverse ways that prokaryotes move. Nat Rev Microbiol, 6, 466–76.

38. Jenal, U., White, J. & Shapiro, L. 1994. Caulobacter flagellar function, but not assembly, requires FliL, a non-polarly localized membrane protein present in all cell types. J Mol Biol, 243, 227–44.

39. Johnson, S., Fong, Y. H., Deme, J. C., Furlong, E. J., Kuhlen, L. & Lea, S. M. 2020. Symmetry mismatch in the MS-ring of the bacterial flagellar rotor explains the structural coordination of secretion and rotation. Nat Microbiol, 5, 966–975.

40. Johnson, S., Furlong, E. J., Deme, J. C., Nord, A. L., Caesar, J. J. E., Chevance, F. F. V., Berry, R. M., Hughes, K. T. & Lea, S. M. 2021. Molecular structure of the intact bacterial flagellar basal body. Nat Microbiol, 6, 712–721.

41. Karimova, G., Pidoux, J., Ullmann, A. & Ladant, D. 1998. A bacterial two-hybrid system based on a reconstituted signal transduction pathway. Proc Natl Acad Sci U S A, 95, 5752–6.

42. Kearns, D. B. 2010. A field guide to bacterial swarming motility. Nat Rev Microbiol, 8, 634–44.

43. Kojima, S., Imada, K., Sakuma, M., Sudo, Y., Kojima, C., Minamino, T., Homma, M. & Namba, K. 2009. Stator assembly and activation mechanism of the flagellar motor by the periplasmic region of MotB. Mol Microbiol, 73, 710–8.

44. Kojima, S., Nonoyama, N., Takekawa, N., Fukuoka, H. & Homma, M. 2011. Mutations targeting the C-terminal domain of FliG can disrupt motor assembly in the Na(+)-driven flagella of Vibrio alginolyticus. J Mol Biol, 414, 62–74.

45. Kunisawa, T. 1995. Identification and chromosomal distribution of DNA sequence segments conserved since divergence of Escherichia coli and Bacillus subtilis. J Mol Evol, 40, 585–93.

46. Kuwajima, G. 1988. Construction of a minimum-size functional flagellin of Escherichia coli. J Bacteriol, 170, 3305–9.

47. Lee, Y. Y. & Belas, R. 2015. Loss of FliL alters Proteus mirabilis surface sensing and temperature-dependent swarming. J Bacteriol, 197, 159–73.

48. Lele, P. P., Hosu, B. G. & Berg, H. C. 2013. Dynamics of mechanosensing in the bacterial flagellar motor. Proc Natl Acad Sci U S A, 110, 11839–44.

49. Lin, T. S., Zhu, S., Kojima, S., Homma, M. & Lo, C. J. 2018. FliL association with flagellar stator in the sodium-driven Vibrio motor characterized by the fluorescent microscopy. Sci Rep, 8, 11172.

50. Liu, X., Roujeinikova, A. & Ottemann, K. M. 2023. FliL Functions in Diverse Microbes to Negatively Modulate Motor Output via Its N-Terminal Region. mBio, e0028323.

51. Lynch, M. J., Levenson, R., Kim, E. A., Sircar, R., Blair, D. F., Dahlquist, F. W. & Crane, B. R. 2017. Co-Folding of a FliF-FliG Split Domain Forms the Basis of the MS:C Ring Interface within the Bacterial Flagellar Motor. Structure, 25, 317–328.

52. Mariconda, S., Wang, Q. & Harshey, R. M. 2006. A mechanical role for the chemotaxis system in swarming motility. Mol Microbiol, 60, 1590–602.

53. Mccarter, L., Hilmen, M. & Silverman, M. 1988. Flagellar dynamometer controls swarmer cell differentiation of V. parahaemolyticus. Cell, 54, 345–51.

54. Mengucci, F., Dardis, C., Mongiardini, E. J., Althabegoiti, M. J., Partridge, J. D., Kojima, S., Homma, M., Quelas, J. I. & Lodeiro, A. R. 2020. Characterization of FliL Proteins in Bradyrhizobium diazoefficiens: Lateral FliL Supports Swimming Motility, and Subpolar FliL Modulates the Lateral Flagellar System. J Bacteriol, 202.

55. Morgenstein, R. M., Szostek, B. & Rather, P. N. 2010. Regulation of gene expression during swarmer cell differentiation in Proteus mirabilis. FEMS Microbiol Rev, 34, 753–63.

56. Motaleb, M. A., Pitzer, J. E., Sultan, S. Z. & Liu, J. 2011. A novel gene inactivation system reveals altered periplasmic flagellar orientation in a Borrelia burgdorferi fliL mutant. J Bacteriol, 193, 3324–31.

57. Mukherjee, S., Bree, A. C., Liu, J., Patrick, J. E., Chien, P. & Kearns, D. B. 2015. Adaptor-mediated Lon proteolysis restricts Bacillus subtilis hyperflagellation. Proc Natl Acad Sci U S A, 112, 250–5.

58. Mukherjee, S. & Kearns, D. B. 2014. The structure and regulation of flagella in Bacillus subtilis. Annu Rev Genet, 48, 319–40.

59. Notredame, C., Higgins, D. G. & Heringa, J. 2000. T-Coffee: A novel method for fast and accurate multiple sequence alignment. J Mol Biol, 302, 205–17.

60. Overhage, J., Bains, M., Brazas, M. D. & Hancock, R. E. 2008. Swarming of Pseudomonas aeruginosa is a complex adaptation leading to increased production of virulence factors and antibiotic resistance. J Bacteriol, 190, 2671–9.

61. Parthasarathy, R. 2012. Rapid, accurate particle tracking by calculation of radial symmetry centers. Nat Methods, 9, 724–6.

62. Partridge, J. D. 2022. Surveying a Swarm: Experimental Techniques To Establish and Examine Bacterial Collective Motion. Appl Environ Microbiol, 88, e0185321.

63. Partridge, J. D. & Harshey, R. M. 2013a. More than motility: Salmonella flagella contribute to overriding friction and facilitating colony hydration during swarming. J Bacteriol, 195, 919–29.

64. Partridge, J. D. & Harshey, R. M. 2013b. Swarming: flexible roaming plans. J Bacteriol, 195, 909–18.

65. Partridge, J. D., Nhu, N. T. Q., Dufour, Y. S. & Harshey, R. M. 2019. Escherichia coli Remodels the Chemotaxis Pathway for Swarming. mBio, 10.

66. Partridge, J. D., Nhu, N. T. Q., Dufour, Y. S. & Harshey, R. M. 2020. Tumble Suppression Is a Conserved Feature of Swarming Motility. mBio, 11.

67. Partridge, J. D., Nieto, V. & Harshey, R. M. 2015. A new player at the flagellar motor: FliL controls both motor output and bias. mBio, 6, e02367.

68. Paul, K., Nieto, V., Carlquist, W. C., Blair, D. F. & Harshey, R. M. 2010. The c-di-GMP binding protein YcgR controls flagellar motor direction and speed to affect chemotaxis by a “backstop brake” mechanism. Mol Cell, 38, 128–39.

69. Pearson, M. M., Rasko, D. A., Smith, S. N. & Mobley, H. L. 2010. Transcriptome of swarming Proteus mirabilis. Infect Immun, 78, 2834–45.

70. Reid, S. W., Leake, M. C., Chandler, J. H., Lo, C. J., Armitage, J. P. & Berry, R. M. 2006. The maximum number of torque-generating units in the flagellar motor of Escherichia coli is at least 11. Proc Natl Acad Sci U S A, 103, 8066–71.

71. Ryu, W. S., Berry, R. M. & Berg, H. C. 2000. Torque-generating units of the flagellar motor of Escherichia coli have a high duty ratio. Nature, 403, 444–7.

72. Sambrook, J. 1989. Molecular cloning: a laboratory manual. Synthetic oligonucleotides.

73. Santiveri, M., Roa-Eguiara, A., Kühne, C., Wadhwa, N., Hu, H., Berg, H. C., Erhardt, M. & Taylor, N. M. I. 2020. Structure and Function of Stator Units of the Bacterial Flagellar Motor. Cell, 183, 244–257.e16.

74. Suaste-Olmos, F., Domenzain, C., Mireles-Rodríguez, J. C., Poggio, S., Osorio, A., Dreyfus, G. & Camarena, L. 2010. The flagellar protein FliL is essential for swimming in Rhodobacter sphaeroides. J Bacteriol, 192, 6230–9.

75. Tachiyama, S., Chan, K. L., Liu, X., Hathroubi, S., Peterson, B., Khan, M. F., Ottemann, K. M., Liu, J. & Roujeinikova, A. 2022. The flagellar motor protein FliL forms a scaffold of circumferentially positioned rings required for stator activation. Proc Natl Acad Sci U S A, 119.

76. Takekawa, N., Isumi, M., Terashima, H., Zhu, S., Nishino, Y., Sakuma, M., Kojima, S., Homma, M. & Imada, K. 2019. Structure of Vibrio FliL, a New Stomatin-like Protein That Assists the Bacterial Flagellar Motor Function. mBio, 10.

77. Tan, J., Zhang, X., Wang, X., Xu, C., Chang, S., Wu, H., Wang, T., Liang, H., Gao, H., Zhou, Y. & Zhu, Y. 2021. Structural basis of assembly and torque transmission of the bacterial flagellar motor. Cell, 184, 2665–2679.e19.

78. Tang, H., Braun, T. F. & Blair, D. F. 1996. Motility protein complexes in the bacterial flagellar motor. J Mol Biol, 261, 209–21.

79. Terasawa, S., Fukuoka, H., Inoue, Y., Sagawa, T., Takahashi, H. & Ishijima, A. 2011. Coordinated reversal of flagellar motors on a single Escherichia coli cell. Biophys J, 100, 2193–200.

80. Terashima, H., Hori, K., Ihara, K., Homma, M. & Kojima, S. 2022. Mutations in the stator protein PomA affect switching of rotational direction in bacterial flagellar motor. Scientific Reports, 12, 2979.

81. Tian, M., Zhang, C., Zhang, R. & Yuan, J. 2021. Collective motion enhances chemotaxis in a two-dimensional bacterial swarm. Biophys J, 120, 1615–1624.

82. Tipping, M. J., Steel, B. C., Delalez, N. J., Berry, R. M. & Armitage, J. P. 2013. Quantification of flagellar motor stator dynamics through in vivo proton-motive force control. Mol Microbiol, 87, 338–47.

83. Togashi, F., Yamaguchi, S., Kihara, M., Aizawa, S. I. & Macnab, R. M. 1997. An extreme clockwise switch bias mutation in fliG of Salmonella typhimurium and its suppression by slow-motile mutations in motA and motB. J Bacteriol, 179, 2994–3003.

84. Toutain, C. M., Zegans, M. E. & O’toole, G. A. 2005. Evidence for two flagellar stators and their role in the motility of Pseudomonas aeruginosa. J Bacteriol, 187, 771–7.

85. Tuson, H. H. & Weibel, D. B. 2013. Bacteria-surface interactions. Soft Matter, 9, 4368–4380.

86. Wadhwa, N. & Berg, H. C. 2022. Bacterial motility: machinery and mechanisms. Nat Rev Microbiol, 20, 161–173.

87. Wang, Q., Mariconda, S., Suzuki, A., Mcclelland, M. & Harshey, R. M. 2006. Uncovering a large set of genes that affect surface motility in Salmonella enterica serovar Typhimurium. J Bacteriol, 188, 7981–4.

88. Wang, Q., Suzuki, A., Mariconda, S., Porwollik, S. & Harshey, R. M. 2005. Sensing wetness: a new role for the bacterial flagellum. Embo j, 24, 2034–42.

89. Waterhouse, A. M., Procter, J. B., Martin, D. M. A., Clamp, M. & Barton, G. J. 2009. Jalview Version 2—a multiple sequence alignment editor and analysis workbench. Bioinformatics, 25, 1189–1191.

90. Wetzel, C., Hu, J., Riethmacher, D., Benckendorff, A., Harder, L., Eilers, A., Moshourab, R., Kozlenkov, A., Labuz, D., Caspani, O., Erdmann, B., Machelska, H., Heppenstall, P. A. & Lewin, G. R. 2007. A stomatin-domain protein essential for touch sensation in the mouse. Nature, 445, 206–9.

91. Xue, C., Lam, K. H., Zhang, H., Sun, K., Lee, S. H., Chen, X. & Au, S. W. N. 2018. Crystal structure of the FliF-FliG complex from Helicobacter pylori yields insight into the assembly of the motor MS-C ring in the bacterial flagellum. J Biol Chem, 293, 2066–2078.

92. Zhou, J., Lloyd, S. A. & Blair, D. F. 1998. Electrostatic interactions between rotor and stator in the bacterial flagellar motor. Proceedings of the National Academy of Sciences, 95, 6436–6441.

93. Zhu, S., Kumar, A., Kojima, S. & Homma, M. 2015. FliL associates with the stator to support torque generation of the sodium-driven polar flagellar motor of Vibrio. Mol Microbiol, 98, 101–10.

## References

94. Amann, E., Ochs, B. & Abel, K. J. 1988. Tightly regulated tac promoter vectors useful for the expression of unfused and fused proteins in Escherichia coli. Gene, 69, 301–15.

95. Calvo, R. A. & Kearns, D. B. 2015. FlgM is secreted by the flagellar export apparatus in Bacillus subtilis. J Bacteriol, 197, 81–91.

96. Datsenko, K. A. & Wanner, B. L. 2000. One-step inactivation of chromosomal genes in Escherichia coli K-12 using PCR products. Proceedings of the National Academy of Sciences, 97, 6640–6645.

97. Guzman, L. M., Belin, D., Carson, M. J. & Beckwith, J. 1995. Tight regulation, modulation, and high-level expression by vectors containing the arabinose PBAD promoter. J Bacteriol, 177, 4121–30.

98. Jumper, J., Evans, R., Pritzel, A., Green, T., Figurnov, M., Ronneberger, O., Tunyasuvunakool, K., Bates, R., Žídek, A., Potapenko, A., Bridgland, A., Meyer, C., Kohl, S. A. A., Ballard, A. J., Cowie, A., Romera-Paredes, B., Nikolov, S., Jain, R., Adler, J., Back, T., Petersen, S., Reiman, D., Clancy, E., Zielinski, M., Steinegger, M., Pacholska, M., Berghammer, T., Bodenstein, S., Silver, D., Vinyals, O., Senior, A. W., Kavukcuoglu, K., Kohli, P. & Hassabis, D. 2021. Highly accurate protein structure prediction with AlphaFold. Nature, 596, 583–589.

99. Kanehisa, M. & Goto, S. 2000. KEGG: kyoto encyclopedia of genes and genomes. Nucleic Acids Res, 28, 27–30.

100. Karimova, G., Pidoux, J., Ullmann, A. & Ladant, D. 1998. A bacterial two-hybrid system based on a reconstituted signal transduction pathway. Proc Natl Acad Sci U S A, 95, 5752–6.

101. Kuwajima, G. 1988. Construction of a minimum-size functional flagellin of Escherichia coli. J Bacteriol, 170, 3305–9.

102. Partridge, J. D., Nieto, V. & Harshey, R. M. 2015. A new player at the flagellar motor: FliL controls both motor output and bias. mBio, 6, e02367.

103. Tang, H., Braun, T. F. & Blair, D. F. 1996. Motility protein complexes in the bacterial flagellar motor. J Mol Biol, 261, 209–21.

104. Varadi, M., Anyango, S., Deshpande, M., Nair, S., Natassia, C., Yordanova, G., Yuan, D., Stroe, O., Wood, G., Laydon, A., Žídek, A., Green, T., Tunyasuvunakool, K., Petersen, S., Jumper, J., Clancy, E., Green, R., Vora, A., Lutfi, M., Figurnov, M., Cowie, A., Hobbs, N., Kohli, P., Kleywegt, G., Birney, E., Hassabis, D. & Velankar, S. 2021. AlphaFold Protein Structure Database: massively expanding the structural coverage of protein-sequence space with high-accuracy models. Nucleic Acids Research, 50, D439–D444.

